# Guidance by followers ensures long-range coordination of cell migration through α-Catenin mechanoperception

**DOI:** 10.1101/2021.04.26.441407

**Authors:** Arthur Boutillon, Sophie Escot, Amélie Elouin, Diego Jahn, Sebastián González-Tirado, Jörn Starruß, Lutz Brusch, Nicolas B. David

**Author notes:** These authors contributed equally (Sophie Escot, Amélie Elouin).

## Abstract

Morphogenesis, wound healing and some cancer metastases depend upon migration of cell collectives that need to be guided to their destination as well as coordinated with other cell movements. During zebrafish gastrulation, extension of the embryonic axis is led by the mesendodermal polster that migrates towards the animal pole, followed by axial mesoderm that undergoes convergence and extension. We here investigate how polster cells are guided towards the animal pole. Using a combination of precise laser ablations, advanced transplants and functional as well as in silico approaches, we establish that each polster cell is oriented by its immediate follower cells. Each cell perceives the migration of followers, through E-Cadherin/α-Catenin mechanotransduction, and aligns with them. Directional information therefore propagates from cell to cell over the whole tissue. Such guidance of migrating cells by followers ensures long-range coordination of movements and developmental robustness.

## Introduction

Cell migration executes and orchestrates key events in development, homeostasis, and disease (Yamada and Sixt, 2019). Apart from a few examples of cells spreading through random migrations (Borrell and Marín, 2006; Pézeron et al., 2008), most cell migration events are precisely guided in vivo, with chemical or physical environmental cues orienting cell movement (Shellard and Mayor, 2020). The past decade has highlighted that many cells do not undertake migration on their own, but are rather influenced by neighbouring cells, in so-called collective migrations (Norden and Lecaudey, 2019; Scarpa and Mayor, 2016; Schumacher, 2019). This is relevant not only to epithelial cells that need to migrate while maintaining close contacts with their neighbours, but also to some mesenchymal cells. The best characterized mesenchymal instance is neural crest cells in Xenopus. In this system, group cohesion is provided by co-attraction, whereby neural crest cells express a chemoattractant to which they themselves respond (Carmona-Fontaine et al., 2011). Concomitantly, cells undergo contact inhibition of locomotion (CIL), whereby contacting cells repolarize away from one another (Carmona-Fontaine et al., 2008; Scarpa et al., 2015). This provides outward polarity for cells in clusters and allows an efficient response to a chemoattractant. While chemoattraction is the best characterized guidance mechanism, some cells rely on mechanical signals for their guidance (Roca-Cusachs et al., 2013). In durotaxis, cells follow gradients in the stiffness of their substrate (Lo et al., 2000; SenGupta et al., 2021; Shellard and Mayor, 2020), a process recently observed in the embryo (Canales Coutiño and Mayor, 2021; Shellard and Mayor, 2021). In addition to following gradients in matrix stiffness, it has been proposed that cells can be guided by physical forces applied at cell-cell contacts (Cai et al., 2014; Tambe et al., 2011). In vitro pulling on Xenopus axial mesoderm cells induces their migration in the opposite direction, a process proposed to guide collective migration in the embryo (Weber et al., 2012).

In addition to being directionally guided, many cell migrations need to be tightly coordinated with other cell movements. This is particularly true during development when many concomitant cell movements shape the forming embryo. In body axis elongation, different cell populations, with different lengthening strategies, need to coordinate their elongation to maintain axis integrity (Bénazéraf et al., 2017; Yang et al., 2002). In fish, mutations slowing some of the axial cells always lead to global slowdown of the axis rather than to a break in the axis (Heisenberg et al., 2000; Topczewski et al., 2001), showing that coordination mechanisms must exist, ensuring axis robustness. Recent work has proposed mechanical interactions as a way to couple movements of different cell populations (Das et al., 2019; Xiong et al., 2020). However, the cellular bases for such interactions, and how long-range coordination can be achieved, remains poorly explored owing to the challenge of both properly imaging and quantifying cell migration in vivo, and of physically altering the cell’s environment to probe the origin and the nature of guidance cues.

Here, we investigate these questions in zebrafish, analysing how the migration of the anterior axial mesendoderm is directed towards the animal pole. At the onset of gastrulation, the first cells to internalize on the dorsal side of the embryo are precursors of the polster (hereafter referred to as polster cells; Solnica-Krezel et al., 1995). From the embryonic organiser, they migrate in straight towards the animal pole, leading the extension of the axis, and are followed by more posterior axial mesodermal cells, including posterior prechordal plate precursors and notochord precursors (Montero et al., 2005; Figure S1A). Although different pathways have been implicated in the migration of polster cells (Blanco et al., 2007; Kai et al., 2008; Montero et al., 2003, 2005; Shimizu et al., 2005; Yamashita et al., 2002, 2004), how polster cells are guided towards the animal pole remains unknown. In particular, loss-of-function of Wnt/Planar Cell Polarity pathway components affects their migration directionality (Heisenberg et al., 2000; Ulrich et al., 2005), suggesting a potential instructive role of the Wnt/PCP pathway in cell guidance. However, ubiquitous optogenetic activation recently demonstrated unambiguously that the Wnt/PCP pathway plays only a permissive role in these cells (Čapek et al., 2019), reopening the question of how they are guided. A few years ago, we demonstrated that migration of polster cells is a collective process: cells require E-Cadherin dependent contacts with their neighbours to perceive directional information and extend protrusions towards the animal pole (Dumortier et al., 2012). Yet the nature of the directional information transmitted at cell contacts and its origin have remained unknown.

In this study, we used complementary approaches, including precise 3D laser ablations and cell transplants, to map the directional information that guides polster cell migration. We identify follower cells as a source, as polster cells need contact with, and animalward movement of the following mesoderm to orient their migration. Cell autonomous inhibition of motility further revealed that polster cells are oriented by the active migration of their immediate followers. Looking at how directional information is conveyed, we find that this is achieved through adherens junctions and the mechanosensitive domain of α-Catenin. These results lead to a model where each cell perceives and is oriented by the migration of its followers, the directional information thus propagating from cell to cell through the tissue. As revealed by numerical simulations, this ‘guidance by followers’ accounts for the long-range coordination of movements of the different cell populations forming the axis, and provides a basis for its developmental robustness.

## Results

### Polster cells do not exhibit Contact Inhibition of Locomotion nor co-attraction

Though they form a group, polster cells appear mesenchymal: all cells form protrusions (Dumortier et al., 2012) and gaps exist between cells (Smutny et al., 2017). Investigating the mechanisms ensuring their collective migration, we first sought to test if polster cells rely on similar processes as those driving neural crest cell migration, namely CIL and co-attraction.

We first tested CIL in polster cells by performing mixing assays (Carmona-Fontaine et al., 2008). Two groups of differently-labelled polster cells were transplanted next to each other at the animal pole of an early gastrula embryo, a region devoid of hypoblast cells that could otherwise interfere with polster cells migration (Figure 1A, Movie S1). The broad mixing observed after 90 minutes argues against the existence of CIL, as illustrated by simulating the assay with cells displaying CIL or not (see Methods; Figure 1A). Second, if CIL was occurring, cells would be expected to change direction upon contact with others (Carmona-Fontaine et al., 2008). For polster cells transplanted at the animal pole, we measured changes in direction upon collision and observed no difference with changes in direction in absence of collision, arguing against a CIL behavior (Figure 1B, Movie S1).

**Figure 1:**
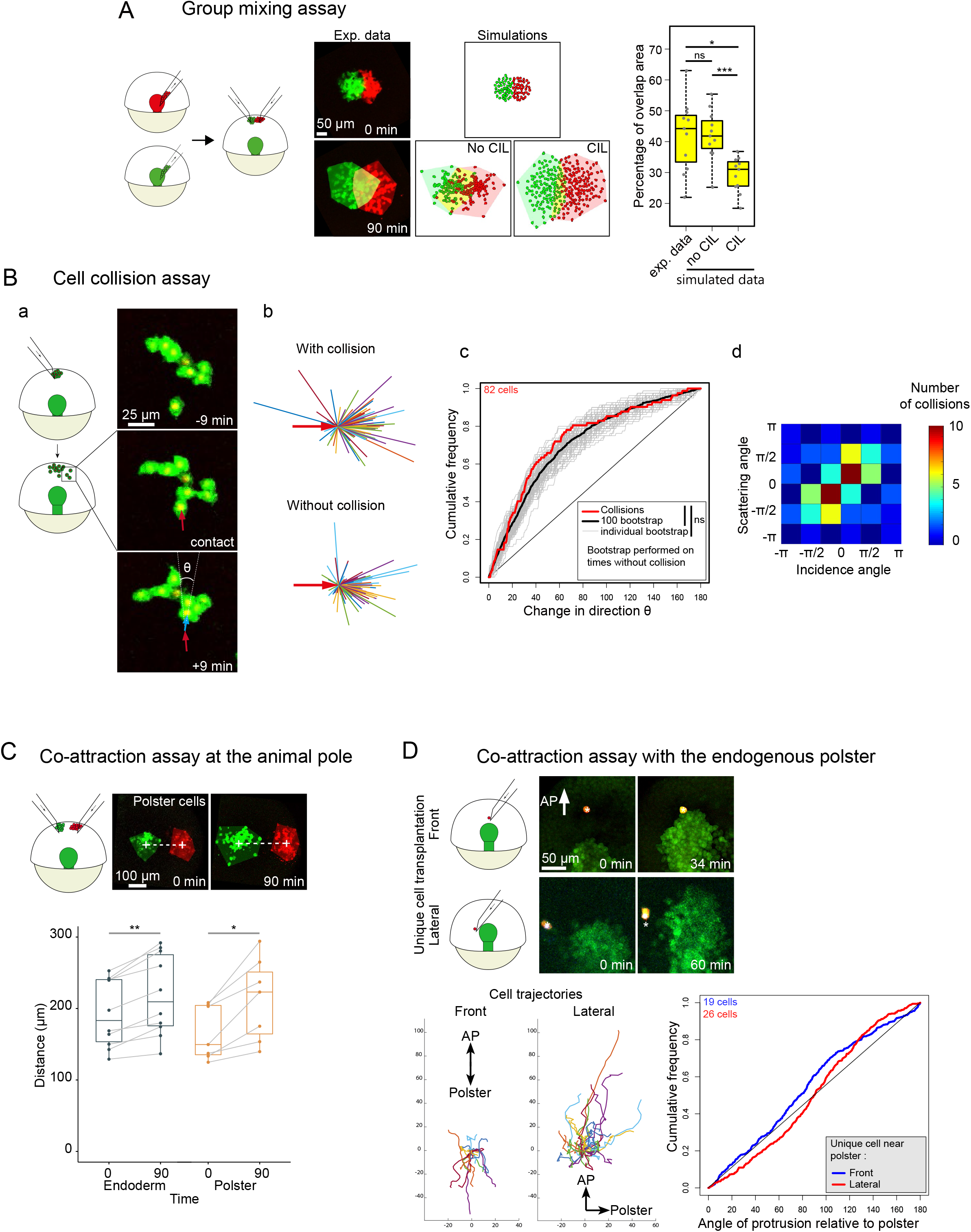
Polster cells do not exhibit CIL nor Coattraction. (A) Mixing of two groups of polster cells, compared to simulations with and without CIL. (B) (a) Collisions of polster cells transplanted at the animal pole. Change in direction (θ) is measured as the angle between the displacement vector before (red arrow) and after (blue arrow) a given time step. (b) Cell trajectories upon collision or not. Initial cell movement is normalized (red arrow), cell movement at the following time step is displayed in various colors. (c) Cumulative frequency of θ upon collision (red) or not (gray: 100 bootstrapped datasets; black: combination of all 100 bootstrapped datasets). (d) Upon collision, the angle between the two cell trajectories after contact (scattering angle) is plotted as a function of the angle between trajectories before contact (incidence angle). CIL would lead to most values being in the top left and bottom right corners, while simple volume exclusion would group values on the diagonal of slope 1 (see D’alessandro et al., 2017). (C) Co-attraction assay of polster or endodermal cells. (D) Unique polster cells transplanted 58±25 μm ahead of or 65±20 μm laterally to the polster. Cell trajectories and orientation of actin-rich protrusions (angle between the direction of the protrusion and the closest polster cell) are displayed for both conditions.

To assess co-attraction, we transplanted two groups of differently-labelled polster cells at the animal pole, separated by some distance, and measured whether they attracted each other (Figure 1C, Movie S1) (Carmona-Fontaine et al., 2011). After 90 min, the distance between centroids had increased by 47±28 μm, a result similar to the one obtained with endodermal cells used as a negative control for co-attraction (Figure 1C; Pézeron et al., 2008). In addition, we observed that groups of cells at the animal pole spread as small clusters or isolated cells; they did not remain compact as would be expected in the case of co-attraction. To further look for signs of co-attraction, we transplanted single polster cells expressing Lifeact-mCherry in front of or laterally to the polster of a host embryo and quantified their movements and protrusion orientations (Figure 1D, Movie S1). Isolated polster cells displayed no preferred movement direction and randomized protrusions, without bias towards the nearby polster. We thus could not detect signs of CIL nor co-attraction in polster cells.

### The directional information guiding the polster does not originate from the polster

We therefore looked for other mechanisms that could guide the collective migration of polster cells. In previous work, we observed that isolated polster cells lack orientation cues, which are restored upon contact with a migrating polster, suggesting that directional cues are present within the polster and transmitted through cell-cell contacts (Dumortier et al., 2012). We sought to map the origin of this directional information. To do so, we developed large 3D ablations, to sever the polster at different positions and identify which regions are required for its oriented migration and where the directional information might come from (see Methods and Boutillon et al., 2021).

We first tested if the first row of polster cells acts as leaders to guide follower cells, a mechanism described in many instances of collective migration (Haeger et al., 2015; Vishwakarma et al., 2020). At 60% epiboly, the front row of cells was ablated (Figure 2A) and the movement of posterior cells was quantified by tracking their H2B-mCherry-labelled nuclei (Figure S2A; Movies S2 and S3). For each cell, we measured both its absolute (aka instantaneous) speed (Figure S2B), and its axial component, in the direction of the animal pole, referred to as axial speed (Figure 2B). Removal of front cells did not affect the absolute speed of following polster cells (Figure S2B), nor their axial speed (Figure 2B), nor the orientation of their protrusions (Figures 2A,C and S2C).

**Figure 2:**
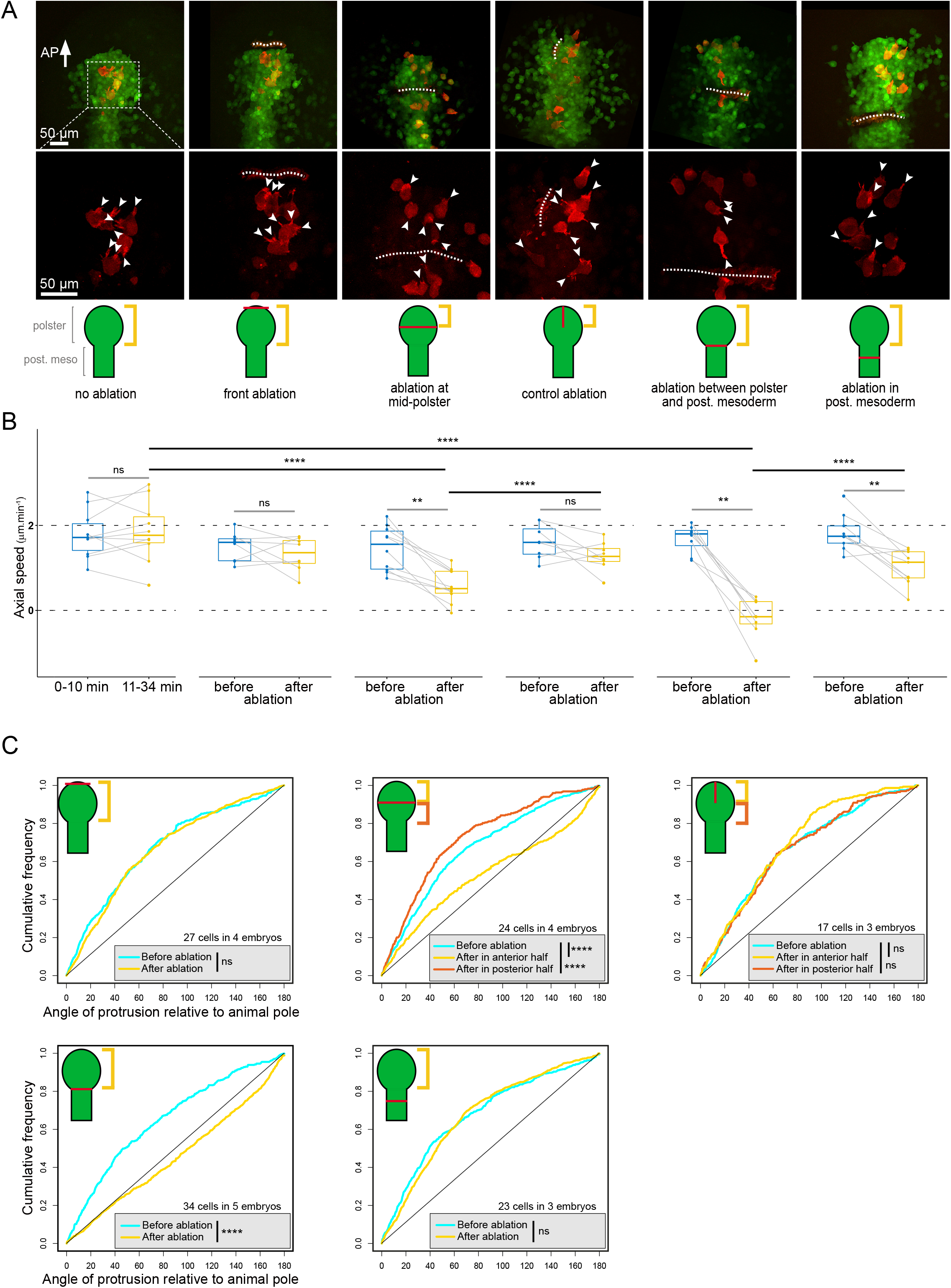
Directional information is not contained in the polster. (A) Laser ablations at varying antero-posterior positions and/or orientations, performed at 60% epiboly. Position of the ablation is indicated by a white dashed line on experimental images, and a red line on schematics; arrowheads mark actin rich protrusions; white arrow indicates direction of the Animal Pole (AP). (B) Axial speed of polster cells, tracked by H2B mCherry labelling of their nuclei. Yellow brackets on schematics indicate the quantified region. n=8 to 10 embryos, 149±11 quantified cells per embryos. (C) Orientation of actin rich protrusions (see also Figure S2C).

To identify the source of directional information, ablations were performed at different antero-posterior positions. First, to isolate the anterior half from the posterior part, middle polster cells were removed by a transversal ablation (Figure 2A, Movie S3). While the absolute speed of anterior cells was not affected by isolation (Figure S2B), their axial speed (animalward motion) decreased dramatically (Figure 2B). As a control for non-specific effects induced by laser ablation, we performed sagittal ablations, parallel to the direction of migration, separating the left and right anterior polster, but leaving each side in contact with the posterior polster (Figure 2A, Movie S3). Such ablations did not significantly reduce axial speed (Figure 2B). Decrease of axial but not absolute speed upon separation of the entire anterior half of the polster suggested that cells exhibited poorer orientation. We tested this by quantifying protrusion orientation of Lifeact-mCherry expressing cells transplanted in the polster before laser ablations. Whereas control sagittal ablations did not affect cell orientation, transversal ablations strongly disrupted protrusion orientation of cells in the isolated anterior polster (Figure 2C). Interestingly, cells in the posterior half were still oriented (Figure 2C) and their axial speed was higher than cells in the anterior part (Figure S2D) suggesting that these cells still perceive a directional information guiding their migration.

To test if the directional information is originating from the posterior polster, we performed similar ablation experiments, this time separating the entire polster from the following axial mesoderm (Figures 2A and S1D, Movie S3). Strikingly, this procedure abolished the animalward movement of polster cells, without affecting their absolute speed (Figures 2B and S2B). Consistent with this loss of direction, the orientation of protrusions was completely lost after ablation (Figure 2C). These experiments reveal that the directional information orienting polster cells is not originating from the polster itself, but rather seems provided by contact with the posterior axial mesoderm.

### Contact with the following axial mesoderm is required for polster oriented migration

To test if the following axial mesoderm is the source of directional information, we performed more posterior ablations, leaving some axial mesoderm in contact with the polster (3.7±1 rows of cells) (Figures 2A and S1D, Movie S3). After ablation, the axial mesoderm at the back of the polster kept elongating (Figure S2E), and, strikingly, polster migration was largely, even though not entirely, restored compared to ablations separating the polster from the following mesoderm (Figure 2B). Consistent with this, cell orientation was also restored (Figure 2C) suggesting that contact between polster and axial mesoderm is necessary for proper orientation and migration of polster cells. This idea is further supported by the observation that, in ablations separating the polster from the following axial mesoderm, the axial mesoderm continued elongating, resulting in wound closure in 24±2 min (n=14 embryos). A few minutes after wound closure, polster migration resumed (Figure S2F) leading to normal development at 24 hpf (Figure S1C).

We sought to confirm these surprising results with a second, independent approach. Using a glass pipette, we removed the polster from *Tg(gsc:GFP)* embryos at 60% epiboly (Figure 3A). In such embryos, posterior axial mesoderm continues elongating. We then transplanted a group of polster cells, with H2B-mCherry labelled nuclei, ahead of the posterior axial mesoderm (Figure 3B, Movie S4). While isolated, these cells spread isotropically. In contrast, after contact with the posterior axial mesoderm, they migrated towards the animal pole (Figure 3C,D). Accordingly, actin-rich protrusions were randomly distributed before contact but became oriented towards the animal pole once the transplanted group was contacted by the posterior axial mesoderm (Figure 3E). In line with laser ablations, these observations demonstrate that the polster requires contact with the following axial mesoderm to orient its migration towards the animal pole.

**Figure 3:**
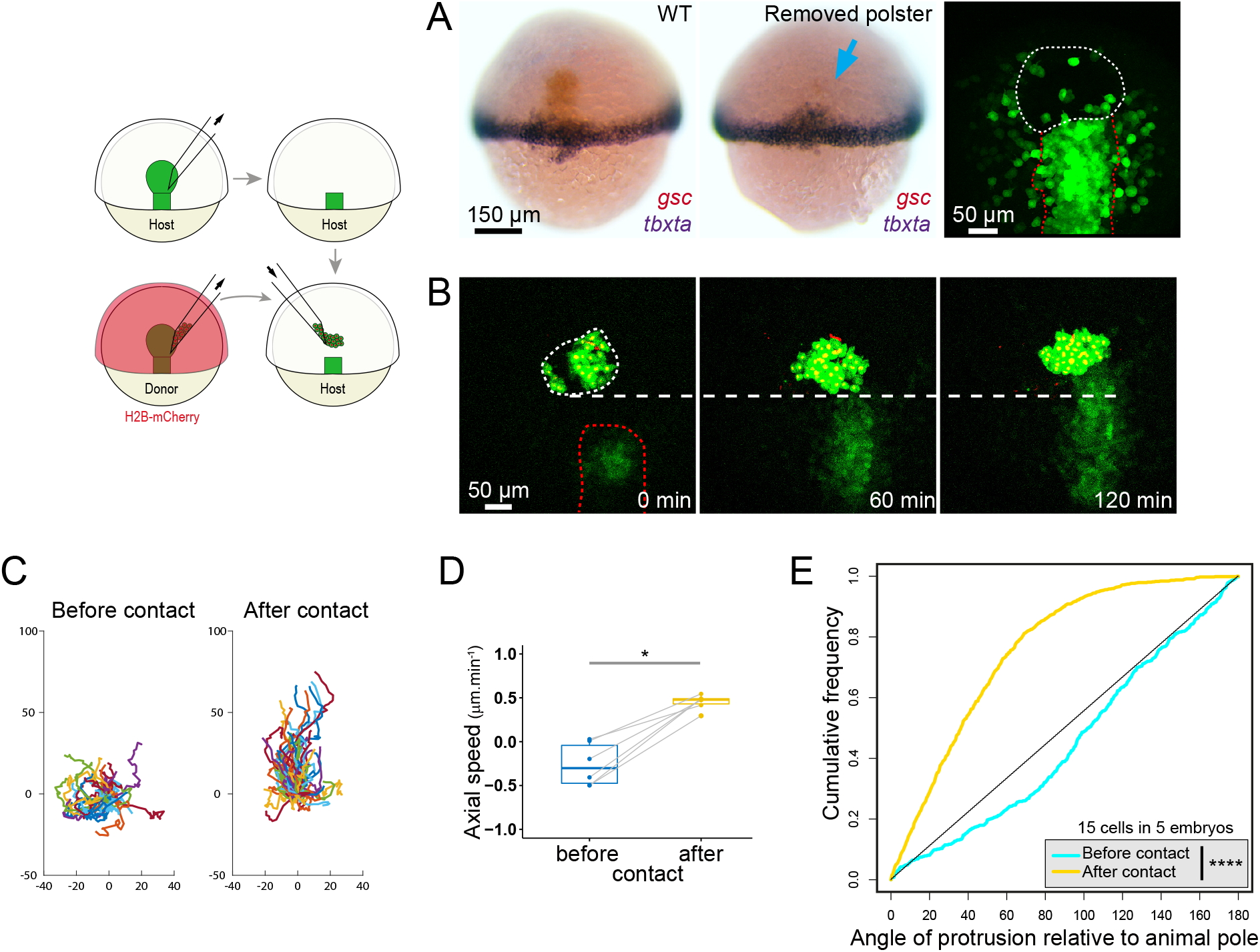
Polster oriented migration requires contact with posterior axial mesoderm. (A) Removal of the polster at 60% epiboly, shown by in situ hybridization for *gsc* (red) and *tbxta* (blue) and fluorescence image in the *Tg(gsc:GFP)* line. Blue arrow and white line mark the former polster position; red lines mark posterior axial mesoderm. (B) Transplantation of 59±51 polster cells, 106±43 μm ahead of the axis. The thin white line delineates transplanted cells; horizontal line marks the initial position of the rear of the transplanted group. (C) Trajectories, (D) axial speed (n=6 embryos) and (E) protrusion orientations of transplanted cells.

### Extension of the following axial mesoderm is required for polster migration orientation

A direct corollary of this need for contact is that extension of the posterior axial mesoderm should be necessary for polster orientation, to maintain contact between the posterior mesoderm and the advancing polster. To test this, we examined the migration of a wild-type polster in front of a defective axis: we genetically slowed axis extension in embryos (see below) and replaced their polster with wild-type polster cells (Figure 4A). As a control, we performed polster replacements between wild-type embryos, and observed no reduction in speed of the polster or of the posterior axial mesoderm compared to untreated embryos (Figure 4B). Inhibiting the non-canonical Wnt-PCP pathway affects axis extension (Čapek et al., 2019; Heisenberg et al., 2000; Ulrich et al., 2005). Consistently, expression of Dsh-DEP+, a dominant negative form of Dsh specifically blocking the PCP pathway (Tada and Smith, 2000), strongly slowed posterior axial mesoderm extension (Figure 4B). Importantly, in Dsh-DEP+ expressing embryos, a transplanted, wild-type, polster showed a similar reduction of its speed towards the animal pole (Figure 4B). A second genetic manipulation that dramatically slows down axial mesoderm extension is expression of Rac1 N17 (Figure 4B), a dominant negative form of the Rac1 small GTPase (DN-Rac1) (Tahinci and Symes, 2003). In DN-Rac1-expressing embryos, the animalward movement of a transplanted, wild-type, polster was essentially abrogated (Figure 4B). In both cases, reduction of speed towards the animal pole appeared due to a loss of cell directionality (Figure 4C), cell protrusions being randomized (Figure 4D). These results demonstrate that the extension of the following axial mesoderm is required for the migration of the polster.

**Figure 4:**
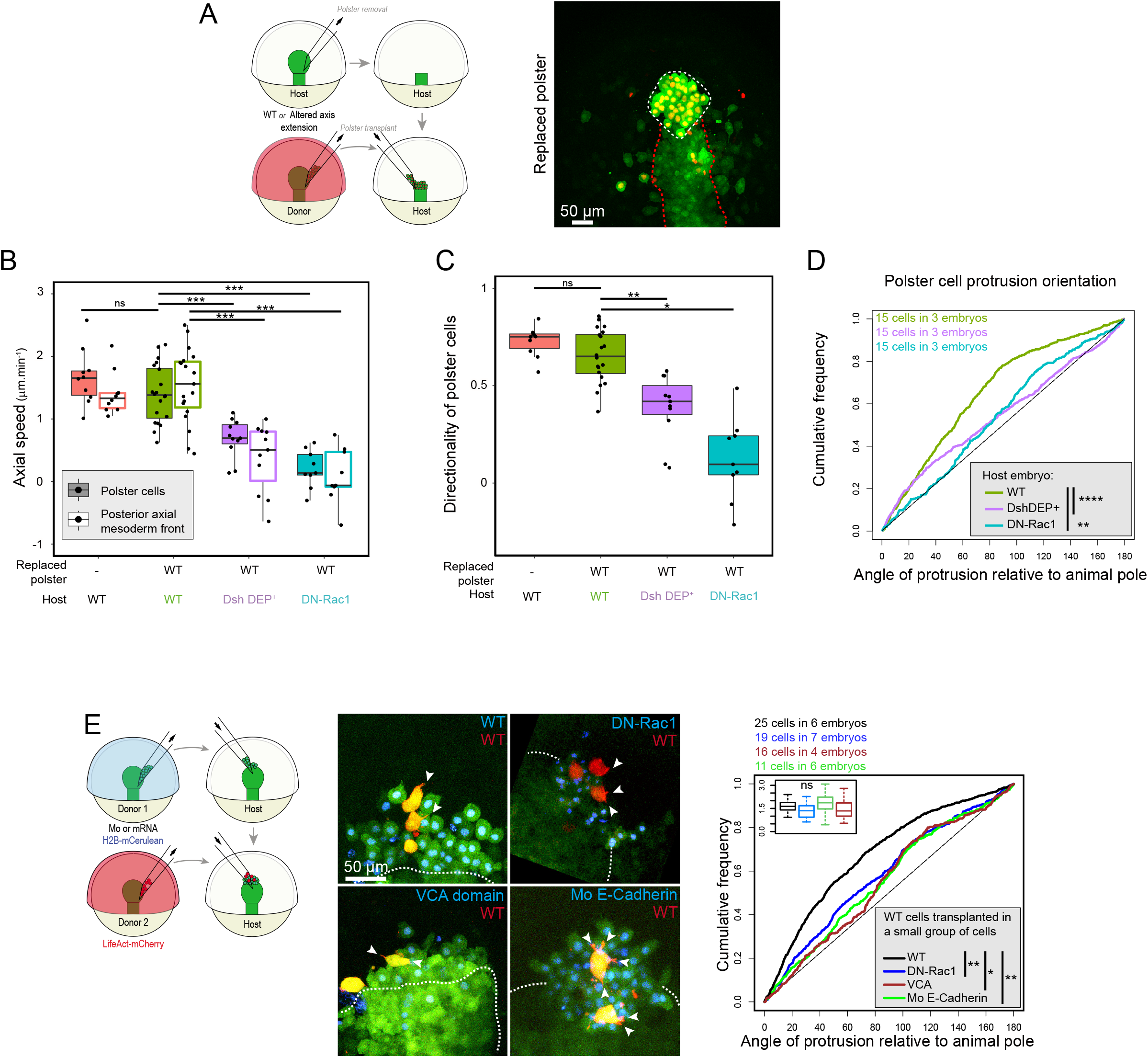
Orientation of polster cells requires active migration of the posterior axial mesoderm. (A) Polster remplacement experiment. White line marks transplanted polster cells and red line marks posterior axial mesoderm. (B) Axial speed of cells in wild-type replaced polster (filled boxes) and of the front of the posterior axial mesoderm (empty boxes) in different conditions. (C-D) Directionality (C) (ratio between axial speed and absolute speed) and protrusion orientation (D) of cells from WT polsters transplanted in different hosts. (E) Protrusion orientation of Lifeact-mCherry expressing WT cells, transplanted among a small group of WT, DN-Rac1, VCA or MO E-Cadherin cells, labelled with H2B mCerulean, in front of a WT polster. Inlay indicates the average number of protrusions per frame of the WT cells, in each condition. White dashed lines mark the endogenous polster of host embryos.

### Active migration of polster cells is required for axis elongation

That the following axial mesoderm extends without polster cells, and that this extension is required for polster migration raised the possibility that the polster is simply passively displaced towards the animal pole by the independently-extending axis. To test whether active migration of polster cells is required, we used DN-Rac1 to inhibit their migration (Dumortier et al., 2012), and transplanted a non-migrating polster into a wild-type embryo. A DN-Rac1-expressing polster replacing that of a wild-type embryo did not move towards the animal pole and blocked the elongation of the axial mesoderm (Figure S3A). Active migration of polster cells is thus required for their movement towards the animal pole. We took advantage of this result to further test the role of axis extension in orienting polster cells, as it provided a way of blocking axis extension without genetically affecting axial cells. Wild-type cells transplanted into a polster expressing DN-Rac1 formed protrusions at the same frequency as in a wild-type replaced polster (Figure S3B inlay). However, in agreement with a requirement for axis extension in orienting polster cells, their protrusions were randomized (Figure S3B).

### Polster cells are oriented by actively migrating followers

Orientation of polster cells by axis extension could result from two mechanisms: one is that axis extension displaces polster cells and this passive displacement is what orients them, the other is that polster cells perceive the active migration of follower cells and use this information to orient their migration. To distinguish between these possibilities, we transplanted a few wild-type cells within a small group of migration defective cells at the front of the polster. Small groups of migration defective cells are displaced by axis extension (Figure S3C,D), so that, in this context, transplanted cells are still displaced but are only in contact with migration defective followers. Migration defective cells were either cells expressing DN-Rac1 or the verprolin, cofilin, acidic (VCA) domain of neural (N)-WASP (Da Costa et al., 2003), both treatments reducing their protrusive activity (Figure S3E). Protrusions of wild-type cells in contact with migration defective cells were as frequent but less oriented than those of cells in a similar small group of wild-type cells (Figure 4E), showing that polster cells require contact with actively migrating followers to become oriented.

If what orients polster cells is the presence of migrating cells at their back, we reasoned that they may not require interactions specifically with the posterior axial mesoderm, and that other cells migrating towards them should be able to orient them. To test this, we transplanted polster cells ahead of the animally migrating lateral mesoderm (Figure S4A, Movie S4, Solnica-Krezel et al., 1995). Before contact, polster cells moved and extended protrusions without a preferred orientation (Figure S4B-D). Upon contact with the lateral mesoderm, they aligned both their migration and protrusions with the movement of the lateral mesoderm. This experiment shows that guidance of polster migration is not specific to posterior axial mesoderm, and can be triggered by another tissue migrating towards polster cell, confirming that having migrating cells at their back is what orients polster cells.

One way that polster cells could be oriented by migrating followers is by perceiving the tensions exerted the follower cells migrating on them. Adherens junctions ensure cell-cell adhesion, can elicit mechanotransduction, and are therefore strong candidates to transmit and perceive tensions. If true, Cadherin should be required both within polster cells to perceive tensions, and in follower cells, to apply tensions. We previously demonstrated (Dumortier et al., 2012), and here confirmed, that E-Cadherin (*cdh1*) knock-down (Figure S5A) cell-autonomously leads to a loss of protrusion orientation, which is rescued by expression of E-Cadherin (Figure 5A-B). At the scale of the entire polster, E-Cadherin knock-down slowed migration towards the animal pole, and reduced cell coherence (Figure S5E). We then tested for a non cell-autonomous requirement for E-Cadherin in neighbouring cells by transplanting wild-type cells within a small E-Cadherin knocked-down cell cluster. The absence of E-Cadherin in neighbours led to a loss of orientation of wild-type cells (Figure 4E). Cadherins are thus required both within polster cells and in their migrating neighbours to orient cell protrusions and migration, consistent with a role in transmitting tensions and their inherent directional information.

**Figure 5:**
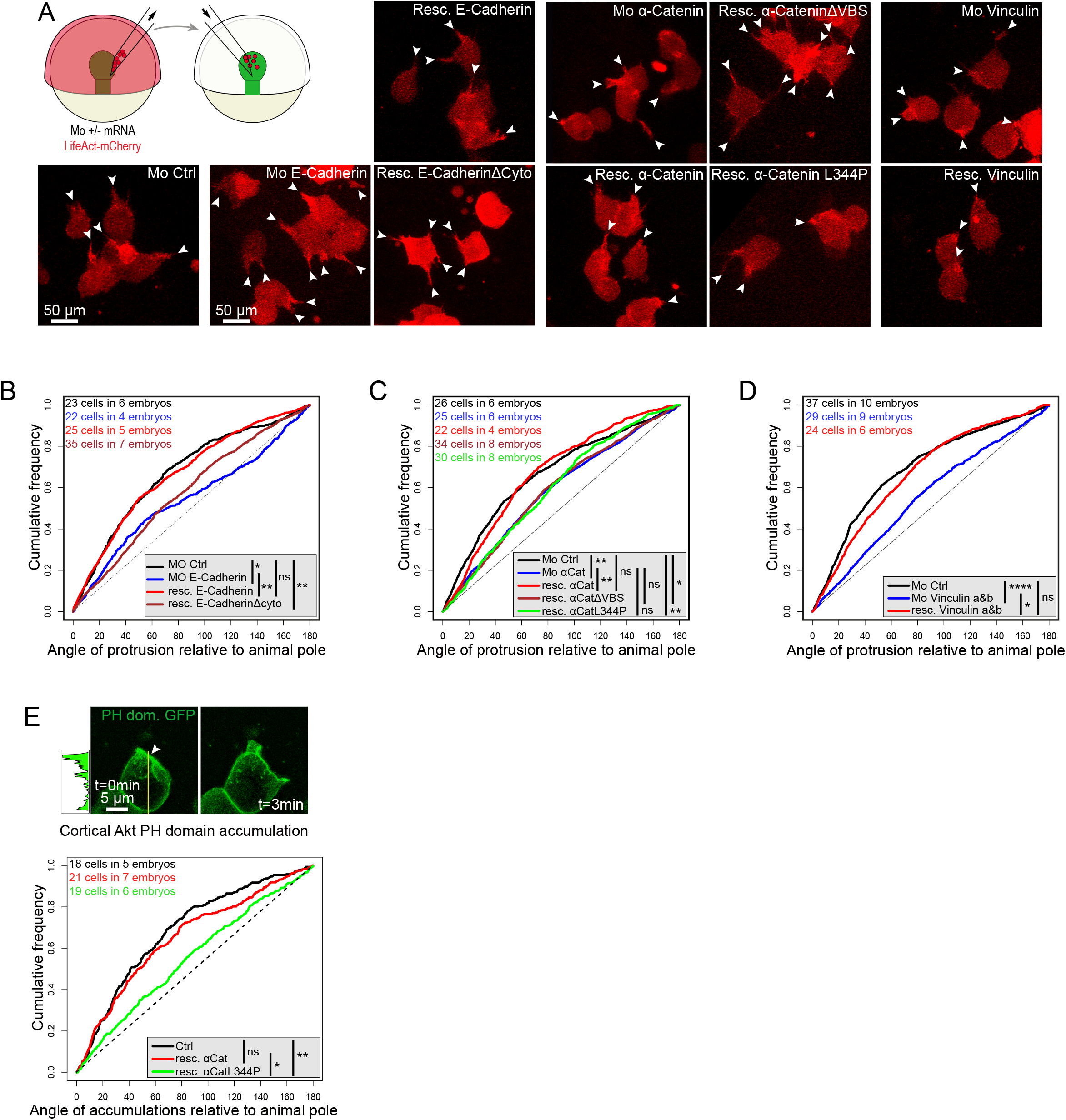
E-Cadherin, α-Catenin mechanosensation and Vinculin are required for polster cell orientation. (A) Protrusions (arrowheads) of different Lifeact-mCherry expressing polster cells, transplanted in a WT polster. (B-D) Protrusion orientations in the different conditions. (E) PH-GFP accumulation at cell front, before formation of a protrusion (arrowhead). Accumultaions are anteriorly biased in control polster cells. This bias is lost in α-Catenin morphant cells expressing the mechano-insensitive form of α-Catenin, but restored upon expression of wild-type α-Catenin.

### Polster cell orientation requires α-Catenin and Vinculin mediated mechanosensation

In some cells, however, Cadherins can influence cell migration without their cytoplasmic domain, in an adhesion and force independent process (Nguyen and Mège, 2016). We ruled this out observing that, contrary to wild-type E-Cadherin, a form of E-Cadherin lacking the intracellular domain (E-CadherinΔcyto, Maître et al., 2012), did not rescue protrusion orientation (Figure 5A,B) of E-cadherin knocked-down cells.

Looking for downstream effectors of E-Cadherin, Plakoglobin was an obvious candidate as, in Xenopus anterior axial mesendoderm, it is recruited to adherens junctions and required for cell orientation in response to tension (Sonavane et al., 2017; Weber et al., 2012). Knocking down both zebrafish orthologs *(jupa* & *jupb)* with morpholinos (Figure S5A) led to cardiac edema and embryonic death at 1 and 3 dpf, as previously described (Figure S5B) (Martin et al., 2009). However, the double knock-down did not affect protrusion orientation of polster cells (Figure S5C).

Another component of adherens junctions, α-Catenin (*ctnna1*), links E-Cadherin to actin and can ensure mechanotransduction (Kobielak et al., 2004; Nieset et al., 1997; Pokutta et al., 2002; Yonemura et al., 2010). Knock-down of α-Catenin (Figure S5A) cell-autonomously reduced protrusion orientation, which could be rescued by co-injection of α-Catenin mRNA, indicating that α-Catenin is required within polster cells for their orientation (Figure 5A,C; Movie S5). At the scale of the entire polster, α-Catenin knock-down slowed migration towards the animal pole, and reduced cell coherence, as E-Cadherin knock-down (Figure S5E). To determine if α-Catenin is required as a link between E-Cadherin and the cytoskeleton or as a mechanosensor, we tried rescuing the knock-down with the α-CateninΔVBS construct, which still links E-Cadherin to actin but lacks mechanosensation (Han et al., 2016; Huveneers et al., 2012; Twiss et al., 2012). As expected, this α-CateninΔVBS partially rescued developmental defects induced by α-Catenin knock-down (Han et al., 2016) (Figure S5D). Yet, it did not restore polster cell orientation (Figure 5A,C and Movie S5) suggesting that the mechanosensory function of α-Catenin is required to orient polster cells. We confirmed these results with the L344P form of α-Catenin, which bears a mutation preventing the tension-dependent recruitment of Vinculin (Seddiki et al., 2018; Figures 5A,C; Movie S5). As the Vinculin binding domain of α-Catenin appeared required for cell orientation, we tested the involvement of Vinculin, knocking down the two zebrafish paralogues *(vcla* and *vclb;* Figure S5A). Consistent with the α-Catenin results, Vinculin is cell-autonomously required for proper cell orientation (Figure 5A,D and Movie S5), and required within the polster for its proper migration (Figure S5E). Formation of polarized protrusions in polster cells has been shown to be dependent on PI3K activation at the cell front (Dumortier et al., 2012; Montero et al., 2003). Using a PH-GFP fusion as a reporter of PI3K activity (Watton and Downward, 1999), we observed that anterior accumulations of PH-GFP were lost in cells lacking α-Catenin mechanosensation (Figure 5E). These results establish that polster cell orientation is driven by mechanotransduction mediated through E-Cadherin, α-Catenin and Vinculin, modulating PI3K activity.

To better understand how follower cells orient cells in front through adherens junction mechanotransduction, we looked at adherens junction localization in polster cells. We observed α-Catenin accumulations at the contact sites between actin-rich protrusions emitted by one cell and the neighbouring cell (Figure 6A). Such accumulations were much more frequent at the rear of cells, contacted by protrusions emitted by the follower cell. These observations suggest that follower cells emit protrusions toward and attach to front cells. These protrusions may impose tension while the follower cell pulls itself forwards. To test this, we assessed the existence of a mechanical tension within protrusions by ablating them and measuring the recoil of the cell. Upon ablation of a protrusion, cells displayed a limited but significant recoil (Figure 6B, Movie S6). As a control, we spotted unattached protrusions, identified as moving protrusions. As expected for unattached protrusions, their ablation led to no recoil. Finally, ablation of protrusions in cells in which adhesion was weakened by E-Cadherin knock-down led to a reduced recoil, consistent with cells applying tensions through adherens junctions.

**Figure 6:**
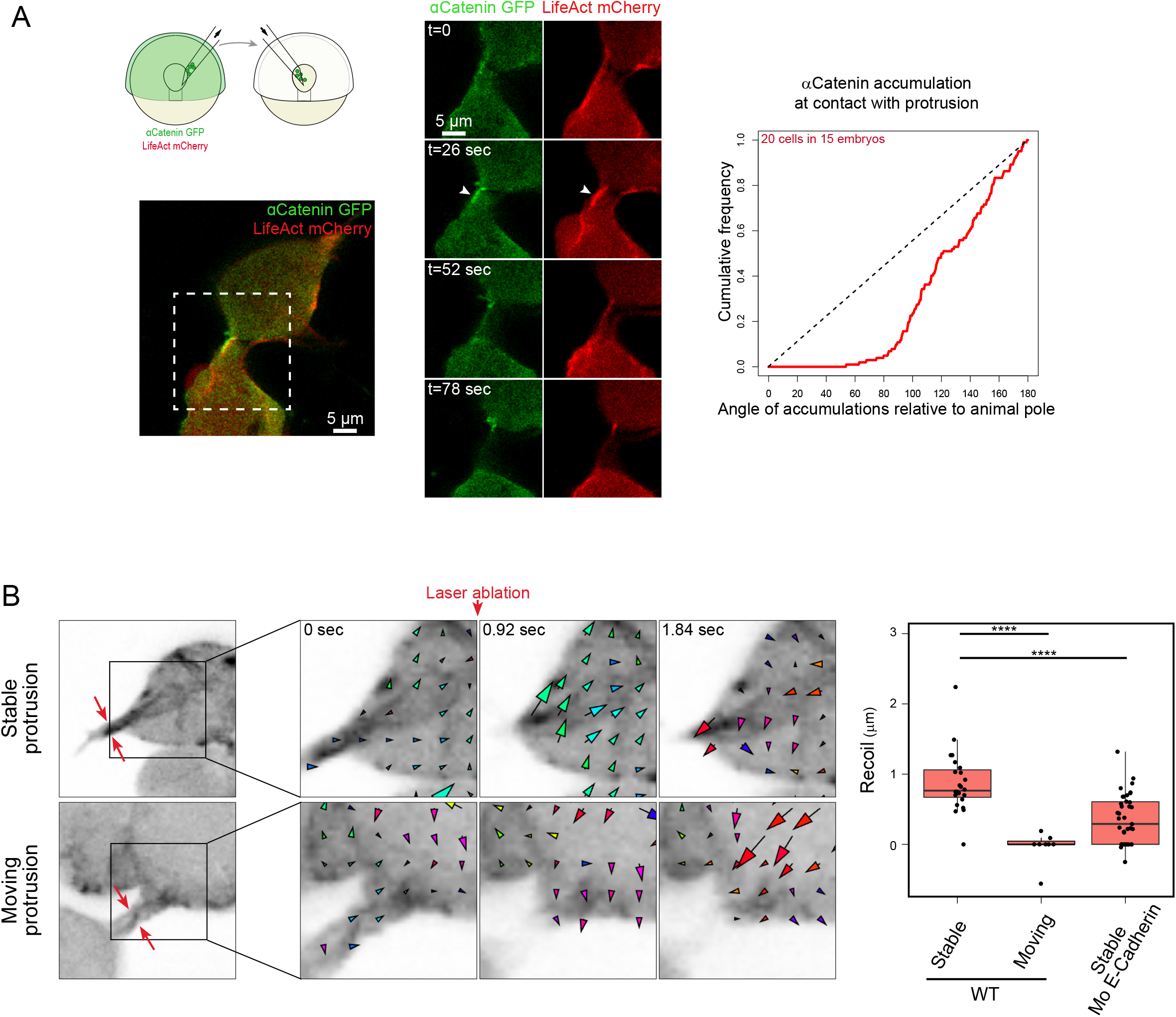
Protrusions attach to cells in front and are under tension. (A) In polster cells, α-Catenin accumulation (arrow head) at the contact site between a protrusion from a follower cell and the back of the cell in front. Positions of accumulations were quantified relative to the direction of the animal pole. (B) Laser ablations of actin rich protrusions in Lifeact-mCherry expressing polster cells. Recoil of the cell was measured immediately after ablation. Ablations were performed on stable protrusions in WT cells (n=24), unattached moving protrusions in WT cells (n=8) and stable protrusions in Mo E-Cadherin cells (n=39).

### In silico simulations reveal the emergence of a robust collective behaviour

Overall, our results suggest that each polster cell is oriented by tensions exerted by actively migrating follower cells, a process we propose to name guidance by followers. To address which statistical properties can emerge from such cell interactions and if this is sufficient to account for the observed collective migration of polster cells, we turned to an in silico approach and used the Cellular Potts Model (Graner and Glazier, 1992) in the modelling and simulation framework Morpheus (Starruß et al., 2014). Briefly, polster cells were given a Run and Tumble behaviour (see Methods and Figure S6; Diz-Muñoz et al., 2016) and a tendency to align with cells migrating towards them. Such a rule proved sufficient to induce the collective migration of polster cells, followed by animalward migrating axial mesodermal cells (Figure 7A, Movie S7). It also correctly reproduced experimental observations upon laser ablations (Movie S7, Figure S7A). In particular, we mimicked the experiments presented in Figure 4A in which the speed of the posterior axial mesoderm is reduced. Consistent with experimental observations, the speed of the simulated polster diminished, even though individual polster cell properties were unchanged (Figure 7B, Movie S7). This reduction of group speed stems from an emergent property of the interacting multicellular system: in simulations, we measured the orientation coherence of polster cells and found it to be linearly dependent on the speed of posterior cells (Figure S7B), so that when axis speed is reduced, polster cells maintain their individual speed, but are less oriented, leading to a reduction of the group speed. We tested this model prediction by measuring cell movement orientation in experimental data and found the same striking correlation between axis speed and polster cell orientation (Figure S7B). Guidance by followers thus coordinates movements of the polster and of the following mesoderm, ensuring axis integrity as an emergent property from cell-cell interactions.

**Figure 7:**
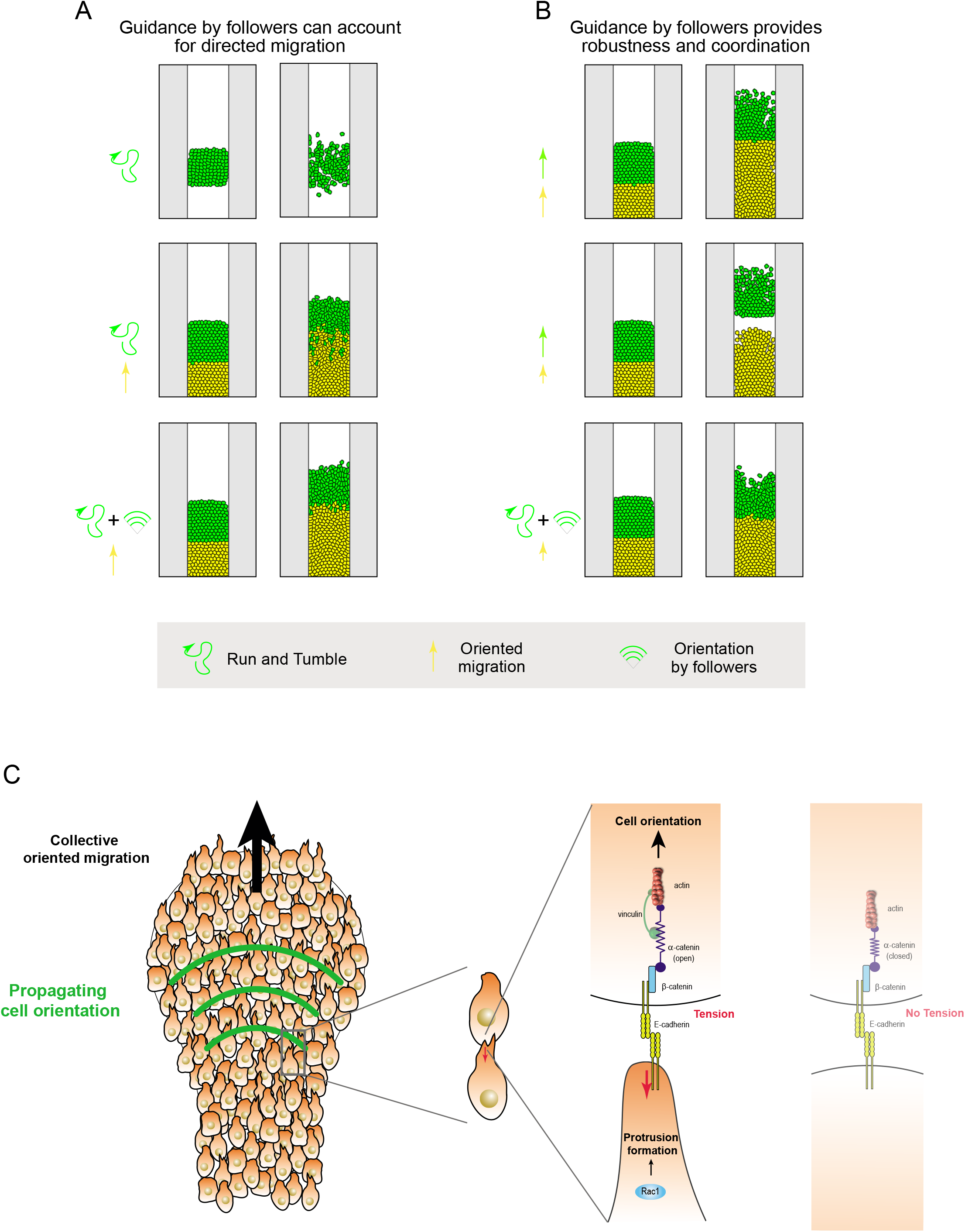
Guidance by followers. (A-B) Cellular Potts models testing different scenarios. (A) Polster cells are given a Run and Tumble behavior, fitted to observations of isolated polster cells (see Figure S6). On their own, polster cells tend to disperse. When followed by axial mesoderm, they progress towards the animal pole, but mix with axial mesoderm. Adding sensitivity to neighbours migrating towards them is sufficient to account for the collective oriented migration of polster cells (and all experimental conditions, see Movie S7). (B) If both polster cells and the following axial mesoderm are given oriented migrations towards the animal pole, differences in their speed will induce axis disruption. On the contrary, when polster cells are guided by followers, polster speed spontaneously adjusts to axis speed. This stems from polster cell orientation being dependent on posterior cell speed (see Figure S7B). (C) Model of guidance of polster cells. Cells perceive, through the mechano-sensitive domain of α-Catenin, the tension generated by the active migration of the following cell, and orient their protrusive activity accordingly. This leads to propagation of the directional information through the entire group.

## Discussion

Precise guidance of migrating cells is required to achieve proper development and morphogenesis. In vitro, many chemical and physical cues can orient cell migration, but it is not clear how such cues can guide cells over long distances in the dynamic environment of the developing embryo. Here, focusing on the forming embryonic axis, we have shown that cells of the polster, the anterior-most cell group, rather than being attracted to their destination by long-range signals, are oriented by the mechano-perception of the follower cells migrating at their back. The directional information thus propagates from cell to cell, creating a robust and self-organizing system. Interestingly, the idea that a mechanical information can propagate and coordinate movements of cells at a distance was recently proposed in two other systems (Das et al., 2019; Xiong et al., 2020). Yet, what generates the mechanical signal, how the mechanical information is perceived, and what cellular properties these signals regulate remained unknown. Here, we established that the mechanical signal is generated by the active migration of follower cells, that it is perceived through the E-Cadherin – α-Catenin – Vinculin pathway, and that it controls cell orientation.

We used laser ablations to map the origin of the directional information guiding polster cells. Ablation of the front row of cells did not affect migration of follower cells, arguing against these cells acting as leaders (Richardson et al., 2016). This is in agreement with previous observations that front and follower cells have similar morphologies and dynamics (Dumortier et al., 2012). We cannot rule out that upon ablation of the front row, follower cells convert into new leaders, but this would have to be done within minutes, as follower cells are not affected, even on short time scales. In addition, if existing, leader cells are not sufficient to drive polster migration, as more posterior ablations, which do not affect the front row, induce a loss of polster cell orientation. Varying the antero-posterior position of ablations, we observed that polster cell orientation is dependent on contact with the following axial mesoderm. In vitro work in Xenopus established that migration of polster cells can be guided by pulling forces applied at the cell rear, leading to a model of collective migration in which the posterior mesoderm would orient migration by acting as a drag (Behrndt and Heisenberg, 2012; Weber et al., 2012). Our observations that posterior axial cells can migrate without polster cells (Figure 3) and that they can displace a group of non-motile cells (Figure S3C) suggest that, in zebrafish, posterior cells are not acting as a drag. On the contrary, our results reveal that it is their active migration which orients polster cell migration. Of note, it is possible that part of the polster movement results from its passive displacement by the progression of the following axial mesoderm. Passive displacement is however not sufficient to account for polster movement, as a non-motile polster remains immobile (Figure S3A), and does not account for polster cell guidance, as polster cells with non-motile followers are not oriented, even though displaced (Figure 4E). Our results rather demonstrate that guidance of one cell is ensured by the migration of its immediate followers. They point to a model where migrating cells emit anteriorward protrusions on which they pull to progress. These protrusions exert tension on cells in front. Tension is perceived through E-Cadherin, α-Catenin and Vinculin mechano-sensation and orients cells (Figure 7C), ensuring cell guidance by followers. In this case, protrusions would serve not only as grapples used to move forward (Lauffenburger and Horwitz, 1996) but also as a means to transmit directional information, a process reminiscent of Drosophila border cells migration (Mishra et al., 2019). Simulations of such cell-cell interactions faithfully reproduced experimental observations, demonstrating that this mechanism is sufficient to propagate directional information across the tissue, and account for the collective behaviour of polster cells.

### How are cells oriented by their neighbours?

Our model suggests that cells perceive tensions exerted at their surface by follower cells and use them to orient actin rich protrusions. We identified the E-Cadherin, α-Catenin, Vinculin pathway as the involved mechanosensor, in line with previous reports showing that stretch induces an α-Catenin change of conformation and Vinculin recruitment (Hoffman and Yap, 2015; Kim et al., 2015; Ladoux et al., 2015). In epithelial cells, this leads to reinforcement of adherens junctions (Jurado et al., 2016) and has been involved in controlling collective epithelial behaviours (Bazellières et al., 2015; Seddiki et al., 2018). How E-Cadherin, α-Catenin and Vinculin control orientation of actin rich protrusions remains to be identified. Our observations suggest this is at least in part mediated through regulation of PI3K activity. Other than that, Merlin appears an attractive candidate, as, in collectively migrating epithelia, it transduces mechanosensation to coordinate Rac1 activity and lamellipodium formation (Das et al., 2015). The Wnt/PCP pathway may also be involved, as it has a well-established role in orienting polster cells (Dumortier et al., 2012; Heisenberg et al., 2000; Ulrich et al., 2005), however this role is permissive only (Čapek et al., 2019; Heisenberg et al., 2000). This raises the question of how PCP is orienting polster cells. It has been proposed that PCP controls the duration of cell contacts (Witzel et al., 2006). In doing so, it could modulate mechanosensation at adherens junctions. Alternatively, in Xenopus ciliated epithelia, planar polarity is determined by mechanical strain (Chien et al., 2015), suggesting that PCP could be set by mechanosensation, and act downstream of mechanosensation in orienting cell migration. In line with this hypothesis, PCP controls compartmentalization of cortical acto-myosin during convergent extension movements in Xenopus (Shindo and Wallingford, 2014).

There is accumulating evidence that mechanosensation at cell-cell contacts has a key role in coordinating many cell behaviours (Hirata et al., 2020; Vassilev et al., 2017; Vishwakarma et al., 2018). However very little is known on how adherens junctions regulate cytoskeleton dynamics and cell migration (Vishwakarma et al., 2020). Unravelling the events downstream of α-Catenin and Vinculin is thus a very exciting avenue for future work, and the zebrafish polster appears as a convenient model system to progress on this line of inquiry.

### Coordination of movements

During development, and during gastrulation in particular, many different cell movements are taking place concomitantly and need to be tightly coordinated to ensure proper morphogenesis. How the different cell populations achieve this coordination remains largely unknown. In body axis elongation, different cell populations, with different lengthening strategies, need to coordinate their elongation to maintain axis integrity (Bénazéraf et al., 2017; Glickman et al., 2003). In fish, it is noticeable that mutants slowing some of the axial cells always lead to global slowdown of the axis, rather than to axis disruption (Heisenberg et al., 2000; Topczewski et al., 2001). This could arise from axial cells all using similar pathways for their migration. Indeed, the Wnt/PCP pathway appears to be required both for the mediolateral intercalations driving posterior axis extension and for the directed migration of polster cells (Roszko et al., 2009). Alternatively, there may be mechanisms ensuring coordination of the two movements. Using transplants of entire polsters, we observed that progression of a wild-type polster is delayed when notochord progression is genetically delayed, revealing the existence of coordination mechanisms ensuring the integrity of the embryonic axis (Figure 4A-D).

Achieving such coordination is not trivial. As illustrated by in silico simulations, systems in which cells have a directed motion towards the animal pole are very sensitive to any difference in speed between polster and more posterior cells (Figure 7B). Adjusting polster speed to axial speed would imply that axial cells instruct polster cells to slow down. On the opposite, guidance by followers spontaneously provides robustness to the system. If posterior cells slow down, they less efficiently orient cells in front, the movement of which is therefore less directed (Figure S7B). This reduces their animalward movement and thus spontaneously adapts their animalward speed to the speed of follower cells (Figure 7B, Movie S7). Guidance by followers, in which the cell-to-cell propagation of directional information orients cell migration, is thus a very simple, yet very effective way of ensuring long-range coordination of cell movements and self-organized guidance. Such mechanical coordination is likely to control morphogenesis in other contexts, in embryonic development, organogenesis or cancer cell migration.

## Limitations of the study

Our work establishes that polster cell guidance is ensured by the animal-ward movement of posterior axial mesodermal cells. This raises the question of how posterior cells are themselves oriented. One intriguing possibility would be that they also rely on guidance by followers, but this remains to be tested. Another limitation is that we performed simulations in 2D while the axial mesoderm is a three-dimensional tissue. It would thus be interesting to verify that a 3D model, closer to tissue architecture, would give similar results. Furthermore, we assumed in simulations that the mesoderm is laterally confined. In the embryo this could be done by the paraxial mesoderm, but this remains to be demonstrated.

## Acknowledgements

We thank F. Rosa for reading the manuscript; F. Graner and W. de Back for initial help with simulations, C. Wyart, J. de Rooij, E. Raz, R.M. Mège, D. Gilmour and M. Breau for fish lines and plasmids; E. Menant for fish care, L. Mellottee for technical help; W. Supatto for help with the PIV analyses; P. Mahou and the Polytechnique Bioimaging Facility for assistance with live imaging on their equipment supported by Région Ile-de-France (interDIM) and Agence Nationale de la Recherche (ANR-11-EQPX-0029 Morphoscope2, ANR-10-INBS-04 France BioImaging). This work was supported by the ANR grants 15-CE13-0016-1, 18-CE13-0024, 20-CE13-0016. S.E. was supported by the European Union’s Horizon 2020 programme under the Marie Skłodowska-Curie grant agreement No 840201. L.B. acknowledges support by the BMBF through FitMultiCell grant 031L0159B. Model simulations for parameter inference were performed on HPC resources granted by the ZIH at TU Dresden.

## Author Contributions

AB and ND conceived experiments, which were performed by AB. SE performed Western Blots and AE helped on localization of α-Catenin. DJ, SGT, JS, LB and ND developed the simulation model, performed the parameter estimation, ran simulations and wrote the corresponding methods section. AB and ND analysed data, wrote the manuscript. LB and ND secured funding.

## Declaration of Interests

The authors declare no competing interests.

## STAR Methods

### RESOURCE AVAILABILITY

#### Lead contact

Further information and requests for resources and reagents should be directed to and will be fulfilled by the lead contact, Nicolas David

#### Materials availability

Plasmids generated in this study were be deposited to Addgene.

#### Data and code availability

- Images used in Figures are available through Mendeley Data.
- The Morpheus model generated during this study was deposited in the public model repository under MorpheusModelID:M0006 (https://identifiers.org/morpheus/M0006). Custom Matlab routines used to process cell tracks and for PIV analysis are available upon request.
- Any additional information required to reanalyze the data reported in this paper is available from the lead contact upon request.

### EXPERIMENTAL MODEL

Zebrafish embryos were obtained by natural spawning of AB, *Tg(tbx16:EGFP)* and *Tg(-1.8gsc:GFP)ml1* adult fishes (Doitsidou et al., 2002; Wells et al., 2011). The *Tg(gsc:GFP)* line labels polster cells, the following axial mesodermal cells, as well as some endodermal cells (Figure S1A, Barone et al., 2017). All animal studies were approved by the Ethical Committee N°59 and the Ministère de l’Education Nationale, de l’Enseignement Supérieur et de la Recherche under the file number APAFIS#15859-2018051710341011v3.

### METHOD DETAILS

#### In Situ Hybridization

Whole-mount colour and fluorescent In Situ Hybridization were performed following standard protocols (Hauptmann and Gerster, 1994) using *goosecoid, tbxta* and *ctslb* probes (Schulte-Merker et al., 1994; Stachel et al., 1993; Thisse et al., 1994).

#### Embryo injection

Translation blocking morpholinos (Gene Tool LLC Philomath) and concentration used were: Vinculin a (5’-CGTCTTGGTATGGAAAACTGGCATC-3’) (0.3 mM), Vinculin b (5’-TGGAAAACCGGCATGATGATCGCTC-3’) (0.3 mM), Jupa (Plakoglobin 1a) (5’-GAGCCTCTCCCATGTGCATTTCCAT-3’) (0.4 mM) (Martin et al., 2009), Jupb (Plakoglobin 1b) (5’-CCTCACTCATTTGCAGTGACATCAC-3’) (0.1 mM), E-Cadherin (5’-TAAATCGCAGCTCTTCCTTCCAACG-3’) (0.3 mM) (Babb and Marrs, 2004), α-Catenin (5’-TAATGCTCGTCATGTTCCAAATTGC-3’) (0.1 mM) (Han et al., 2016), Sox32 (5’-CAGGGAGCATCCGGTCGAGATACAT-3’) (0.3 mM) (Dickmeis et al., 2001), and standard control (5’-CCTCTTACCTCAGTTACAATTTATA-3’) (0.1 to 0.3 mM).

Capped sense mRNA were synthesized from pCS2+ or pSP64T constructs with mMessage mMachine SP6 kit (Thermo Fischer). Constructs and concentrations used were: *histone2B-mCherry* (30 to 50 ng/μl), *histone2B-mCerulean* (30 to 50 ng/μl), *lifeact-mCherry* (30 to 50 ng/μl), *acvr1ba\** (0.6ng/μl), *dshDep^+^* (75 ng/μl), *rac1N17* (2 or 10 nl/μl), *Zf cdh1-GFP* (60 ng/μl), *Zf cdh1-Δcyto-GFP* (60 ng/μl), *Zf ctnna1* (30 ng/μl), *Zf ctnna1-ΔVBS* (30 ng/μl), *Zf ctnna1L344P* (30 ng/μl), *Zf ctnna1-GFP* (30 ng/μl), *Zf GFP-vcla* (25ng/μl), *Zf GFP-vclb* (25ng/μl) and *PH-GFP* (75ng/μl).

To label and/or affect the whole embryo, 5 nl were injected at the one-cell stage. For donor embryos for cell transplantation, 1.5 nl were injected in one cell at the four-cell stage.

#### Cell transplantation and microsurgery

Cell transplantations were performed as described in (Boutillon et al., 2018). Cells transplanted within the polster were taken from the shield of a *Tg(gsc:GFP)* donor and transplanted to the shield of a *Tg(gsc:GFP)* host at 6 hpf. Identity of transplanted cells was then assessed by their GFP expression. Cells transplanted out of the polster (animal pole, lateral side, ahead of the polster) were taken from donor embryos injected, in one cell out of four, with *acvr1ba\** mRNA and Sox32 morpholino, so as to impose a polster identity (Dumortier et al., 2012). For single cell transplant, donor embryos were dissociated at shield stage in Ringer’s without calcium solution prior to transplantation. Removal of the polster was performed in *Tg(gsc:GFP)* embryos, by aspiration with a large homemade glass pipette. The polster was identified on morphological criteria, confirmed by *in situ* hybridization against *ctslb,* a marker for polster identity, and *gsc*, a marker for prechordal plate (Figures 3A and S1).

#### Image analysis and cell movement quantifications

Cell movements were quantified by tracking cell nuclei, labelled with H2B-mCherry, using IMARIS (Bitplane). Tracks were then processed using custom-made Matlab (Math Works) routines as described in (Dumortier et al., 2012), to compute absolute speed, axial speed, directionality and coherence. Coherence is actually measured as a coherence^-1^: for each cell, it is the change in distance to its neighbors (defined as cells within a 30-μm radius), normalized by the cell net displacement over the considered time interval. Posterior axial mesoderm elongation was quantified by tracking migration of cells at its front. Actin-rich protrusions were quantified on Lifeact-mCherry expressing cells.

#### Quantification of protrusion orientation

Quantifications of actin-rich protrusions were performed on Lifeact-mCherry expressing cells. A few polster were transplanted into unlabelled hosts, providing scarce labelling, which allows precise quantification of cell protrusions and measurement of their orientations (Boutillon et al., 2018). Using ImageJ (FIJI), a line was manually drawn from the centroid of every labelled cell to the basis of each of its protrusions. Angle of protrusion relative to the direction of polster migration was then measured to obtain the protrusion angular distribution for each cell. Protrusion frequency of each cell was finally obtained by averaging the number of protrusions at each frame.

#### Mixing and co-attraction assays

For mixing assay, two differently labelled groups of polster cells (red and green) were transplanted adjacently, at the animal pole of an early gastrula embryo (shield stage). Overlap, highlighted in yellow, is measured after 90 min, for experimental and simulated data. Mixing of the two groups was measured after 90 minutes as the normalized area of overlap between the two cell populations. Simulated cells display migration characteristics similar to polster cells (see also Fig S6), with or without CIL behavior.

For co-attraction assay at the animal pole, two differently labelled groups of polster or endodermal cells were transplanted at the animal pole of an early gastrula embryo (shield stage), initially 166±38 and 197±44 μm apart respectively. Distance between group centroids was measured at 0 and 90 min.

#### Embryo imaging

Imaging of embryos for protrusion quantification was done on an inverted TCS SP8 confocal microscope (Leica) equipped with environmental chamber (Life Imaging Services) at 28°C using a HC PL APO 40x/1.10 W CS2 objective (Leica). Imaging of embryos for cell migration quantification was done under an upright TriM Scope II (La Vision Biotech) two-photon microscope equipped with an environmental chamber (okolab) at 28°C and a XLPLN25XWMP2 (Olympus) 25x water immersion objective or on the inverted TCS SP8 microscope (Leica) using a HCX PL Fluotar 10x/0.3 objective (Leica). Injected embryos were mounted in 0.2% agarose in embryo medium between 60% and 70% epiboly (6.5-7.5 hpf). Embryos were imaged between 30 and 60 minutes, every one to three minutes.

#### Laser ablation

Laser ablation experiments were performed under the TriM Scope II microscope (La Vision Biotech) equipped with a femtosecond Mai Tai HP DeepSee laser (Spectra Physics) and an Insight DeepSee (Spectra Physics) laser (Boutillon et al., 2021).

For large ablations, severing the polster, embryos were imaged every minute for 10 to 15 minutes prior to ablation. GFP was excited by the Mai Tai laser set to 920 nm wavelength and mCherry by the Insight laser set to 1160 nm. Ablations were performed with the Mai Tai laser at 820 nm and exit power at 0.3 mW. Such an exit power allowed efficient ablation with very good axial confinement. The region to be ablated was defined as an XY ROI, and selectively illuminated using an EOM. To perform 3D ablations, laser treatment was performed on different focal planes, separated by 10 to 15 microns, starting with deeper planes. To compensate for the loss of energy in deeper planes, the number of treatment repeats was modulated with depth. Efficiency of the ablation was assessed by the absence of GFP fluorescence and the presence of cellular debris, and later confirmed by observation of locally modified cell behaviour. Embryos were imaged for 30 to 40 minutes following ablation. The polster was identified on morphological criteria and distance to the front, confirmed by *in situ* hybridization against *ctslb,* a marker for polster identity (Figure S1D). The use of non-linear optics provided sufficient axial resolution not to affect the yolk cell underneath, nor the ectoderm above (Figure S1B). Treated embryos could develop until at least 24 hours post fertilization (hpf) and presented only a slight delay compared to controls, suggesting that the laser treatment was not harmful (Figure S1C).

For protrusion ablations, mCherry was excited by the Insight laser set to 1160 nm, ablations were performed with the Mai Tai laser at 820 nm and exit power at 0.4 mW. Imaging was limited to 50 μm wide areas, to reduce the time interval between frames to 0.3 s. 8 to 10 images were recorded before ablation of a 5x×7 microns area across the protrusion. Depending on cell depth in the embryo, ablation was performed over 1 to 3 time frames. Imaging was restarted immediately after ablation, over 50 time frames. Recoil was measured both manually on FIJI and through PIV analysis. Both approaches led to similar results. PIV analysis was based on MatPIV, adapted to fluorescent images (addition of a threshold on the signal-to-noise ratio, to limit analysis to regions of the image containing signal).

#### Western blots

Embryos were collected at mid-gastrulation (8 hours post fertilization), dechororinated and deyolked manually. Five embryos were pooled and lysed in RIPA and 1X proteinase inhibitor solution. The lysate was boiled in Laemmli SDS-PAGE sample buffer for 5 min. Sample separation by SDS-PAGE electrophoresis was performed using Mini-Protean 10% TGX Precast gels (Bio-Rad). After electrophoresis, protein gels were blotted to Nitrocellulose membranes using the Trans-Blot Turbo RTA Transfer System and the Trans-Blot Turbo Transfert Buffer (Bio Rad). Membrane was blocked for 1 h at room temperature in Odyssey Blocking Buffer (Li-Cor). Primary and secondary antibodies were diluted in Blocking Solution. Antibodies against α-Catenin, α-Tubulin, E-cadherin, Plakoglobin were used at 1/4000, antibody against Vinculin was used at 1/500, secondary antibodies were used at 1/5000. The membrane was incubated in primary antibody over night at 4°C, washed five times for 10 min with PBS with 0,1% Tween 20 and incubated with the secondary antibody for 60 min at room temperature before being washed, as with primary antibody. Images were acquired using a ChemiDoc MP (Bio-Rad).

#### Model simulations

##### Model description

To model cell motility and cell-cell interactions, we chose a Cellular Potts Model (CPM) since the CPM allows for arbitrary cell shapes, spatially resolved cell-cell contact interfaces and stochasticity in cell movement (Graner and Glazier, 1992). Multiple modelling and simulation frameworks for CPM exist including Chaste (Mirams et al., 2013; Pitt-Francis et al., 2009), CompuCell3D (Swat et al., 2012) and Morpheus (Starruß et al., 2014) which are free, open-source software. We have chosen Morpheus because of its user-friendly interface and its transparent separation of the solver code from the computational model description in the domain-specific language MorpheusML. The model description file was deposited in the public model repository under MorpheusModelID:M0006 (https://identifiers.org/morpheus/M0006) which renders our multicellular simulations readily reproducible and extensible following the FAIR principles.

The simulations were performed on an elongated spatial domain with 500 x1500 grid nodes of a two-dimensional hexagonal lattice with periodic boundary conditions. Left- and right-flanking static obstacles left a central channel of 200 nodes width for the cells to migrate into. These obstacles were used to mimic lateral confinement by paraxial mesoderm. The spatial unit is chosen as 1μm per grid interval and the temporal unit as 1 min per time step. Monte Carlo step duration was chosen as 0.1 min to allow thousands of potential updates per lattice node during the simulated time span. Cell shape is controlled by a target area of experimentally measured 326 μm^2^ (average of 360 experimental measures) and a target circumference taken from the isoareal circle. Both constraints enter the Hamiltonian with equal Lagrange multipliers of 1 (Graner and Glazier, 1992). Axial mesoderm cells (yellow in simulations) are given a directed motion targeted at the animal pole, the speed of which is modulated by varying the strength parameter of the Directed Motion plugin in Morpheus (Starruß et al., 2014). To characterize the cell-autonomous behavior of polster cells (without guidance by followers), we transplanted wild-type polster cells to the animal pole of wild-type embryos (or to the margin of MZ*oep* embryos), and tracked the migration of isolated cells. Tracks displayed alternating phases of relatively straight migration and phases of slowed and poorly directed movement, in agreement with a previous report, demonstrating that mesendodermal cells display run-and-tumbling (Diz-Muñoz et al., 2016). We therefore implemented a run-and-tumble motility, with uniform reorientation probability of the target direction, a non-dimensional scaling factor “Run_duration_adjustment” of the Gamma-distributed probabilistic waiting times for reorientation events, and a tunable Lagrange multiplier “motion_strength” that scales motion speed. The two parameter values for Run_duration_adjustment and motion_strength were estimated from experimental data of single cell trajectories (see parameter estimation). We found good quantitative agreement with the experimental data, see Fig. S6C-D. To test whether a simpler model could also match our experimental data, we implemented the biased random walk model (Codling et al., 2008; Weiss, 1994) with uniform choice of direction at each step and an optional directional bias, hence two fit parameters. This biased random walk model was fit against the same data as done for the Run-and-Tumble model but resulted in a fivefold remaining error at best fit and was thus ruled out.

In addition to the Run-and-Tumble motility, mechanical orientation of polster cells was simulated using the PyMapper plugin. At fixed time steps of 1 min, for each cell, neighbours are detected on 50 membrane points. For each neighbour, the angle between its velocity vector and the direction towards the considered cell is computed. If below a threshold “max_angle” (i.e. neighbour migrating towards the considered cell), the velocity vector of the neighbour is used as the new direction of the considered cell in the Directed Motion plugin. In case of several migrating neighbours, the direction vector is an average of their velocity vectors, weighed by the size of cell-cell contacts. In most simulations, 400 cells are initialized. An initial phase of 20 min without motility is used to equilibrate cell shapes and cell packing (not shown on the movies). Based on their antero-posterior position, cells are then given an identity, and the corresponding motility properties. The main parameters were: target cell area (A_0_) set to 326 μm^2^ based on experimental measurements, target cell circumference (C_0_) set to √(4πA_0_), motion strength of polster cells (λ_1_) at 0,5 fitted to single cell data, mean run time of polster cells (T_1_) at 0,76*2,5 min fitted to single cell data, advection velocity of polster cells (v_1_) at 1,42 μm/min fitter to single cell data, maximum angle (α_max_) at π/6 fitted to collective behaviour.

For simulations with Contact Inhibition of Locomotion, identical parameters were used (Monte Carlo step duration, target area and circumference) and polster cells were given the same Run and Tumble behaviour. Instead of adding mechanical orientation, a CIL behaviour was added. Briefly, a membrane property is used to detect contact with neighbouring cells. The vector between the cell center and the contact point is measured and the opposite vector is added to the current cell direction, with a tunable Lagrange multiplier “cil_strength”. Simulations were performed on a square domain with 1000 x1000 grid nodes of a two-dimensional hexagonal lattice with periodic boundary conditions. Two groups of 40 cells were initialized, their centres 120 μm apart.

##### Parameter estimation

In order to fit the baseline cell motility parameters to experimentally observed single cell trajectory data, we define a distance function between the observed and simulated summary statistics.

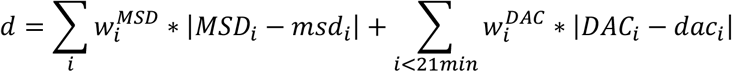

Here, MSD is the mean square displacement and DAC the direction autocorrelation function (Gorelik and Gautreau, 2014). Capital variables represent the experimental measurements, small letters represent the model observables. We calculate the sum of weighted (see below) differences between the ensemble (n~200) means of MSD and DAC at the time points i ∈ {0min, 3min, 6min, 9min, 12min, 15min, 18min, 21min}. We optimize this distance function *d* employing the FitMultiCell software (https://fitmulticell.gitlab.io), which is a free, open-source Python tool embedding stochastic, multi-cellular Morpheus simulations in the highly parallel and unbiased Approximate Bayesian Computation - Sequential Monte Carlo (ABC-SMC) algorithm implemented by the computational framework pyABC (Klinger et al., 2018; Schälte and Hasenauer, 2020). FitMultiCell concurrently evaluates the model and distance measure *d* for trial parameter sets drawn from the evolving probability distribution across the search space, started from a uniform prior distribution. The distance measure *d* is then minimized over successive epochs by only accepting parameter sets with *d* below a gradually decreasing acceptance threshold ε. The pyABC meta-parameters for parameter sample number was set to 200 accepted trial parameter sets per epoch. The computations were run on the high performance computing cluster of ZIH at TU Dresden with 4 CPUs used per task, 2.5 GB/core. The approximate run time for 10 epochs was 20 hrs.

The weights wi were chosen adaptively in pyABC to account for the different scales of MSD and DAC and two sets of optimization epochs were concatenated. First, a uniform prior distribution in the broad interval [0.01, 10] was chosen for each fit parameter and the adaptive-weight scheme of pyABC readjusted the wi after each of 10 epochs. To avoid convergence problems for later epochs due to fluctuating weights, we used the posterior distributions of this first set of epochs as the prior for the second set of epochs, i.e. Run_duration_adjustment: [0.05, 4], motion_strength: [0.1, 2], advection_velocity: [0.05, 3]. The weights wi of the second set of epochs were again adaptively adjusted by pyABC but just initially and then kept fixed for the remaining 12 epochs. Convergence was judged by arriving at the plateau of the acceptance threshold ε in the ABC-SMC algorithm, see Figure S4. The following point estimate for the fitted model parameters and their confidence intervals were obtained (Figure S6A):

Run_duration_adjustment: 0.76, CI: [0.18, 1.95]
motion_strength: 0.50, CI: [0.28, 1.00]
advection_velocity: 1.42, CI: [0.24, 2.20]

The parameter advection_velocity was used to overlay a uniform translation onto all cells, capturing potential drag forces by the overlying ectoderm. Such common translation reproduces the experimentally observed baseline of 20% in DAC (Figure S6D).

#### Illustrations

Images were processed with FIJI. Figures were assembled with Adobe InDesign, movies with Adobe Premiere Pro.

### QUANTIFICATION AND STATISTICAL ANALYSIS

All statistical analyses were performed in R (R project). Cell migration absolute and axial speed were averaged over cells and embryos, and compared using Wilcoxon tests. When relevant, paired Wilcoxon tests were performed, and are indicated on the corresponding boxplots by grey lines joining paired data. Protrusion angle distributions and frequency were compared using linear mixed models taking into account the fact that measurements are not independent (several measurements for each cell, several cells for each embryo). In all figures, ns: p.value ?0.05; *: p<0.05; **: p<0.01; ***: p<10-3; ****: p<10-4.

To serve as controls for collision data (see Figure 1B), 100 bootstrapped datasets were generated by randomly picking, for 80 cells, one time-step where the cell moves freely (no collision) and measuring the angle between incident and efferent vectors (26±13 available times per cell). Each of these bootstrapped datasets, along with the combination of all, were compared to the angle of deflection upon collision using a Kolmogorov-Smirnov test. Only one bootstrapped dataset is statistically different, which is less than expected by chance with an α risk of 5%.

## Supplemental Information

**Movie S1**: **Polster cells do not exhibit CIL or CoA behaviour**, related to Figure 1. Mixing assay: two differently labelled groups of polster cells were transplanted side by side at the animal pole of a host embryo. Their overlap (yellow) was measured after 90 min. Green is GFP, red is Lifeact-mCherry. Collision assay: example of a collision between polster cells transplanted at the animal pole of a host embryo. Nuclei, labelled with Histone2B-mCherry were tracked (white ellipsoid). Two nearby groups: two differently labelled groups of polster cells were transplanted 160 μm apart, to see if they attract each other. Areas covered by each group are highlighted in green and red at times 0 and 90 min. Green is GFP, red is Lifeact-mCherry. Trajectories before and after contact are highlighted, respectively in cyan and yellow. One cell ahead of the polster: one polster cell, labelled with Lifeact-mCherry (red), was transplanted ahead of the polster (green). Scale bar is 50 μm for all four movies.

**Movie S2**: **Four-dimensional tracking of polster nuclei**, related to Figure 2. Nuclei of a *Tg(gsc:GFP)* embryo were labelled with Histone2B-mCherry. Z-stacks were acquired every minute. Nuclei of polster cells, identified by GFP expression and morphological criteria, are highlighted in magenta and 3D-tracked in time. Animal pole is to the top. Scale bar is 50μm.

**Movie S3: Laser ablations**, related to Figure 2. Maximum projections of z-stacks acquired every minute in gastrulating *Tg(gsc:GFP)* embryos. Embryos were first imaged for 10 minutes. A laser ablation was then performed at the location indicated in each panel by the red bar on the movie and on the small schematic. Embryos were then imaged for 40 minutes. Animal pole is to the top. Scale bar is 50 μm.

**Movie S4**: **Polster cells are oriented by actively migrating followers**, related to Figures 3 and S4. Contact with posterior axial mesoderm drives polster cell migration: the polster of an unlabelled *Tg(gsc:GFP)* host was removed, while the following axial mesoderm was left intact (dim green). Polster cells from a *Tg(gsc:GFP)* donor (bright green), labelled with Histone2B-mCherry (red nuclei), were transplanted ahead of the remaining axial mesoderm. Z-stacks were acquired every 2-minute, maximum projections are shown here. Before contact with the extending axial mesoderm, transplanted polster cells do not display directional migration. After contact, the group of polster cells migrates in the same direction as the extending axial mesoderm. Contact with the lateral mesoderm can orient polster cell migration: polster cells from a *Tg(gsc:GFP)* donor (bright green), labelled with Histone2B-mCherry (red nuclei), were transplanted ahead of the lateral mesoderm of a *Tg(tbx16:GFP)* embryo (dim green). Z-stacks were acquired every 3-minute, maximum projections are shown here. Trajectories of representative cells are highlighted. Before contact with the lateral mesoderm, polster cells tend to spread (cyan tracks), while after contact they align with the lateral mesoderm (yellow tracks). Animal pole is to the top. Scale bar is 50 μm.

**Movie S5**: **α-Catenin and Vinculin mediated mechanosensation is required for polster cell orientation**, related to Figure 6. Polster cells injected with morpholino (Mo) or morpholino and mRNA (Resc.) and labelled with Lifeact-mCherry were transplanted in the polster of wild-type embryos and acquired in 3D over time. Maximum projections are shown. Time-interval between frames is, from left to right, top to bottom: 90 s, 60 s, 120 s, 60 s, 60 s, 120 s and 90 s. Animal pole (AP) is to the top. Scale bar is 20 μm.

**Movie S6**: **Ablation of a stable and of a moving protrusion in Lifeact-mCherry expressing polster cell**, related to Figure 6. Left: imaging of the Lifeact-mCherry signal; right: PIV analysis.

**Movie S7**: **Simulations of polster migration**, related to Figure 7. ‘Mechanical orientation (guidance by followers) can account for directed migration of the polster’: movies of the three scenarios presented on Figure 7A. ‘Simulations reproduce experimental observations’: polster cells are given a Run and Tumble behaviour and mechanical sensitivity to neighbours migrating towards them (guidance by followers). When isolated cells, polster cells tend to disperse, an equivalent experimental condition (polster cells transplanted to the animal pole of a host embryo) is presented for comparison. Simulating laser ablations at the front does not affect polster migration, as observed (Figure 2, Movie S3). Simulating laser ablation at the interface between polster and following mesoderm halts progression of the polster till wound healing, as observed (Figure 2, Movie S3). ‘Each cell responding to all its neighbours cannot account for experimental observations’: other simple rules, like correlating the movement of one cell to the movement of any of its neighbours, as observed in confluent epithelia (Poujade et al., 2007) for instance, could not reproduce experimental. It can account for the directed migration of the polster (left movie) but cannot account for the behaviour of an isolated polster (center movie) or for the behaviour after laser ablation at the interface between the polster and the following mesoderm (right movie). ‘Mechanical orientation (guidance by followers) provides robustness and coordination’: movies of the three scenarios presented on Figure 7B. Coordination of the progression of the polster and of the following mesoderm can be achieved by giving both tissues a directed migration (left movie). However, such a system is very sensitive to differences in cell speed (center movie). On the opposite, when polster cells are mechanically oriented, polster speed adjusts to the speed of the following mesoderm, ensuring axis continuity. This stems from polster cells being less oriented when followed by slow axial cells (see Figure S5B).

## Notes

### Competing Interest Statement

The authors have declared no competing interest.

