## Supplemental Figures for "Guidance by followers ensures long-range coordination of cell migration through α-Catenin mechanoperception"

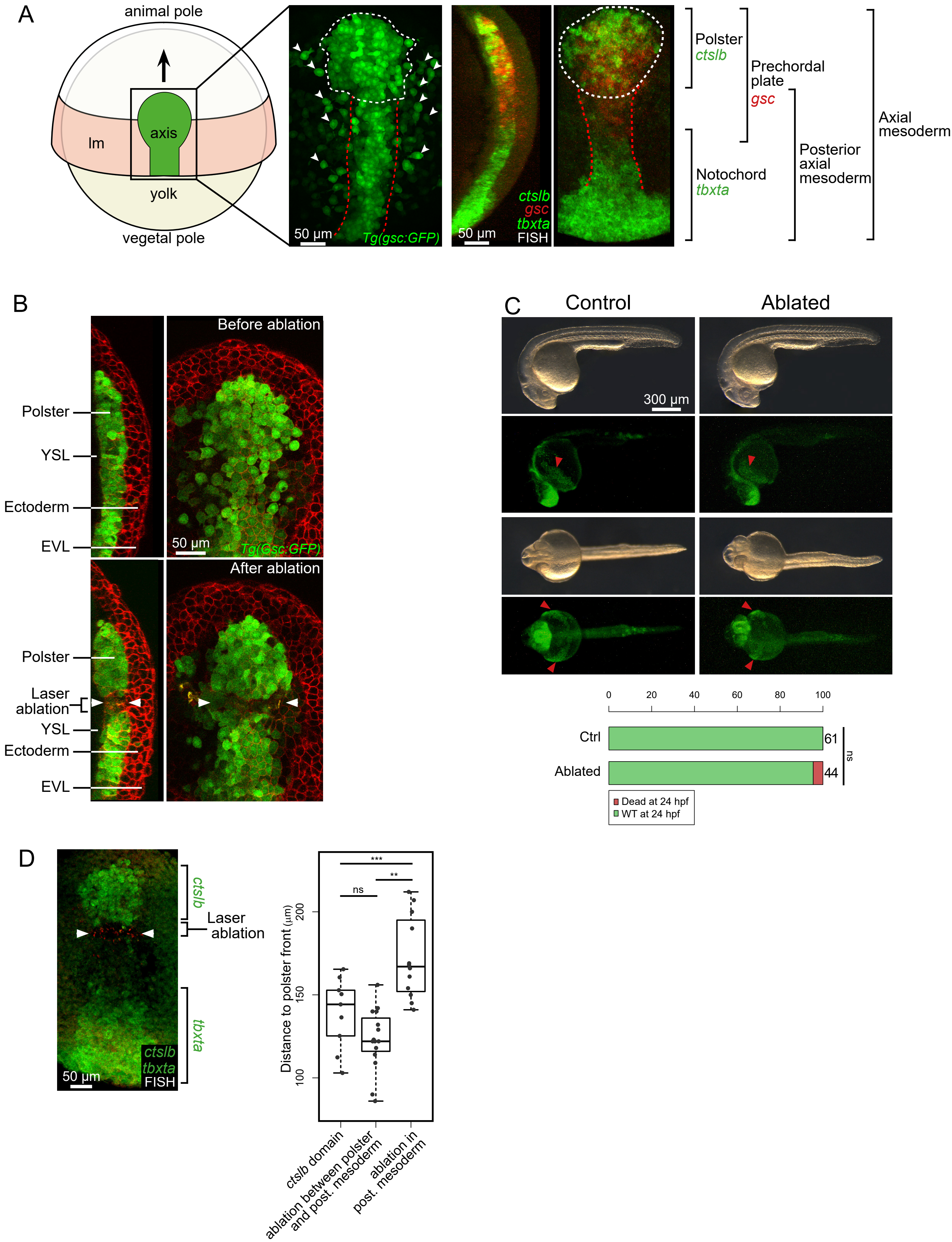


**Supplemental Figure 1. Situation of polster cells and laser ablations,** related to Figure 2.
(A) Dorsal view of a gastrulating embryo at 75% epiboly; lm: lateral mesoderm; black arrow marks the direc-

tion of polster migration. Dorsal view of a Tg(gsc:GFP) embryo where axial mesoderm is labelled in green, along with some endodermal cells; white line delineates the polster; red lines mark the posterior axial mesoderm; arrowheads point to some endodermal cells. Fluorescent in situ hybridization (FISH) of the gastrulating zebrafish axial mesoderm, sagittal and dorsal views. Polster precursors expressing ctslb and notochord precursors expressing tbxta appear in green, prechordal plate progenitors expressing gsc appear in red.
(B) Representative Tg(gsc:GFP) embryo before and after laser ablation, here between the polster and the posterior mesoderm. Sagittal and dorsal views. Membranes are labelled in red by expression of mCherry-CAAX. Ablation is located between white arrowheads.
(C) Morphology and survival of control and ablated Tg(gsc:GFP) embryos at 24hpf. The polster derivative, the hatching gland, is indicated by red arrowheads.
(D) Fluorescent in situ hybridization for ctslb and tbxta on a representative embryo ablated at the interface between the polster and the posterior axial mesoderm. Position of the ablation is visible through red autofluorescent debris. Position relative to the front of the polster of the back of the ctslb domain, of the position of ablations between the polster and posterior mesoderm, and of ablations within the posterior mesoderm.

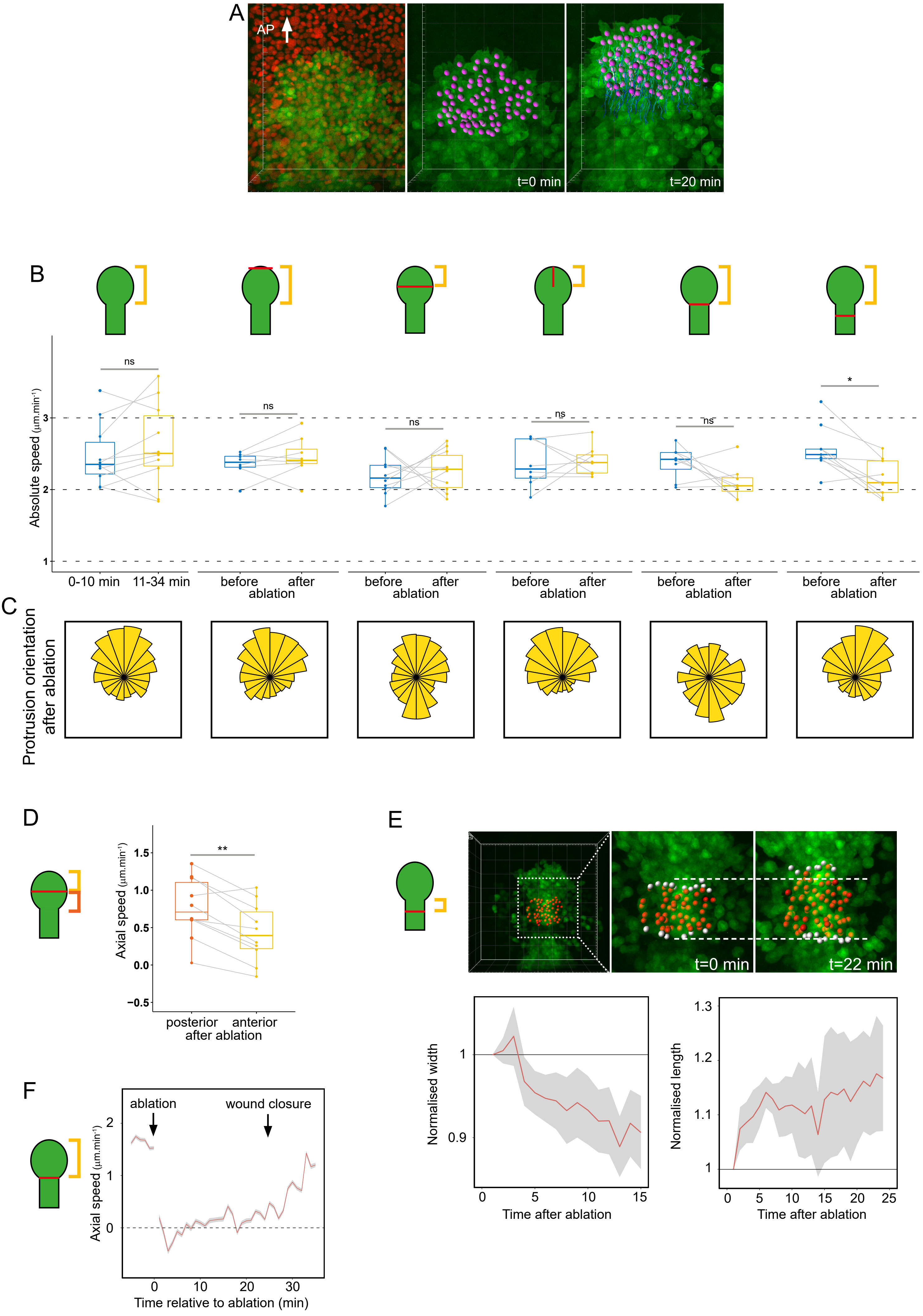
A
AP
t=0 min
t=20 min
B
ns
ns
ns
ns
ns
*
Absolute speed (µm.min⁻¹)
0-10 min
11-34 min
before
after
ablation
C
Protrusion orientation after ablation
D
**
Axial speed (µm.min⁻¹)
posterior
anterior
after ablation
E
t=0 min
t=22 min
Normalised width
Time after ablation
Normalised length
Time after ablation
F
ablation
wound closure
Axial speed (µm.min⁻¹)
Time relative to ablation (min)

**Supplemental Figure 2. Migration of polster cells after laser ablations,** related to Figure 2.
(A) 3D projections showing polster and posterior axial mesoderm migration, in a Tg(gsc:GFP) embryo expressing H2B-mCherry (red). Nuclei belonging to the polster are highlighted in magenta and tracked over time (see Movie S2). AP: animal pole.
(B) Absolute speed of polster cells in control and ablated embryos, corresponding to the axial speeds presented in Fig. 2B.
(C) Radial plots of the orientation of protrusions of polster cells, after ablation, in the area indicated by the bracket. Corresponds to the yellow cumulative plots in Fig. 2C.
(D) Axial speed of the anterior and posterior parts of the polster after ablation in its middle.
(E) Convergence and extension of the posterior axial mesoderm immediately at the back of the polster, after its separation from the rest of the posterior axial mesoderm. White spots mark anterior- and posterior-most isolated posterior mesoderm nuclei. Normalized length and width of the posterior mesoderm were measured over time after ablation.
(F) Axial speed of polster cells in embryos ablated between the polster and the posterior mesoderm, as a function of time (n=8 embryos). The moment of the ablation and the moment of wound healing are indicated.
(B-F) Schematics indicate the position of ablation; brackets indicate the regions quantified after ablation.
(B,D) Gray bars indicate paired statistical test on embryos before and after ablation.

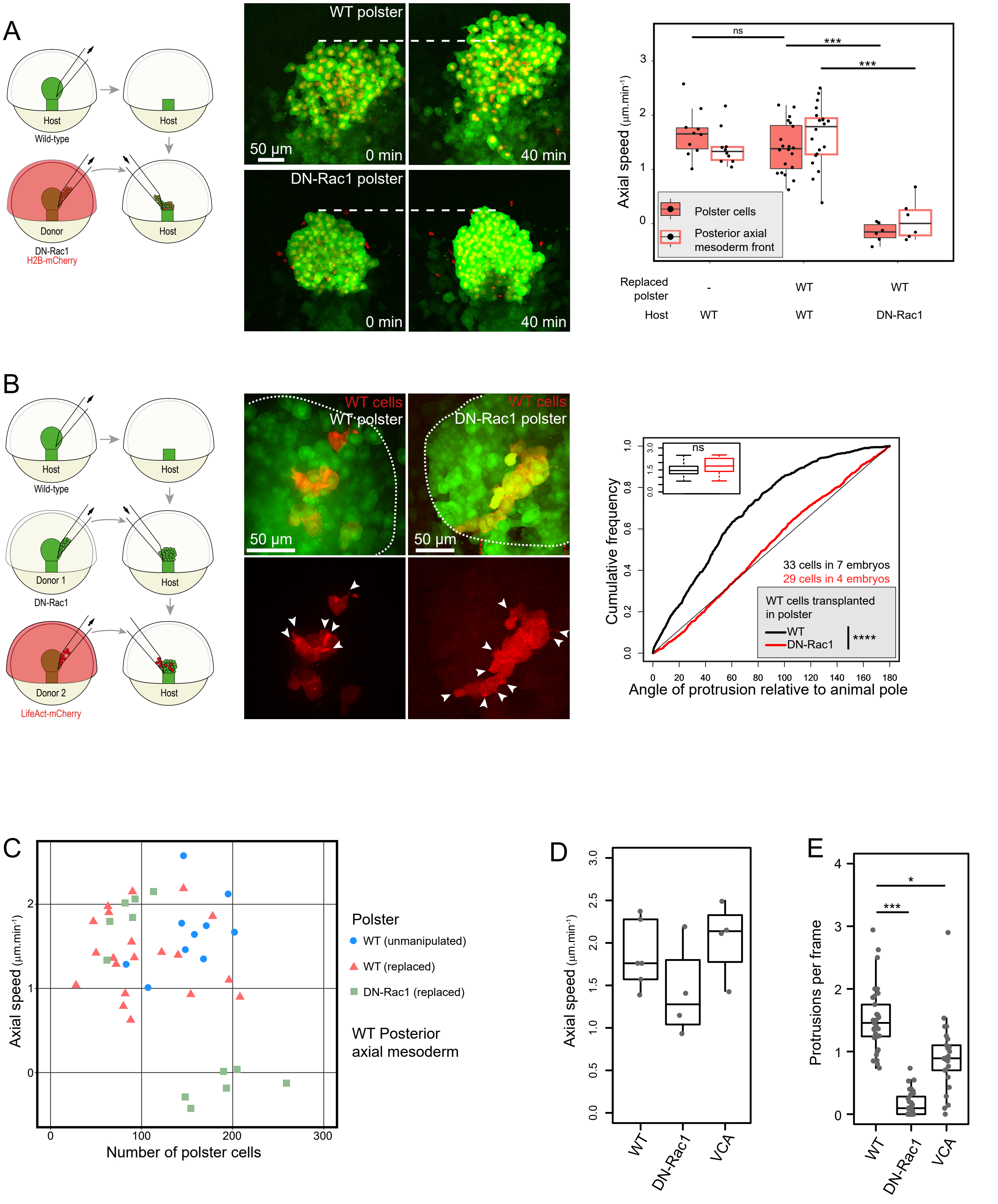


**Supplemental Figure 3. Movement of migration defective polster cells or wild-type polster cells with migration defective neighbours,** related to Figure 4.

(A) Polster replacement experiments. Contrary to a wild-type polster, a DN-Rac1 polster does not migrate towards the animal pole. Lines mark the initial position of the polster front.

(B) Protrusion orientation of wild-type (WT) polster cells in WT (33 cells in 7 embryos) or DN-Rac1 (29 cells in 4 embryos) polsters. Inlay indicates the average number of protrusions per WT transplanted cell and per frame, for both conditions.

(C) Polster replacement experiments, with WT or DN-Rac1 polsters, of varying size. DN-Rac1 polsters the size of an endogenous polster (WT unmanipulated) do not move towards the animal pole. Small DN-Rac1

polsters are however displaced towards the animal pole by the extending axis.
(D) Axial speed of WT, DN-Rac1 or VCA expressing polster cells, transplanted in front of a WT polster (see Figure 4E).
(E) Quantification of protrusions in WT, DN-Rac1 or VCA expressing polster cells.

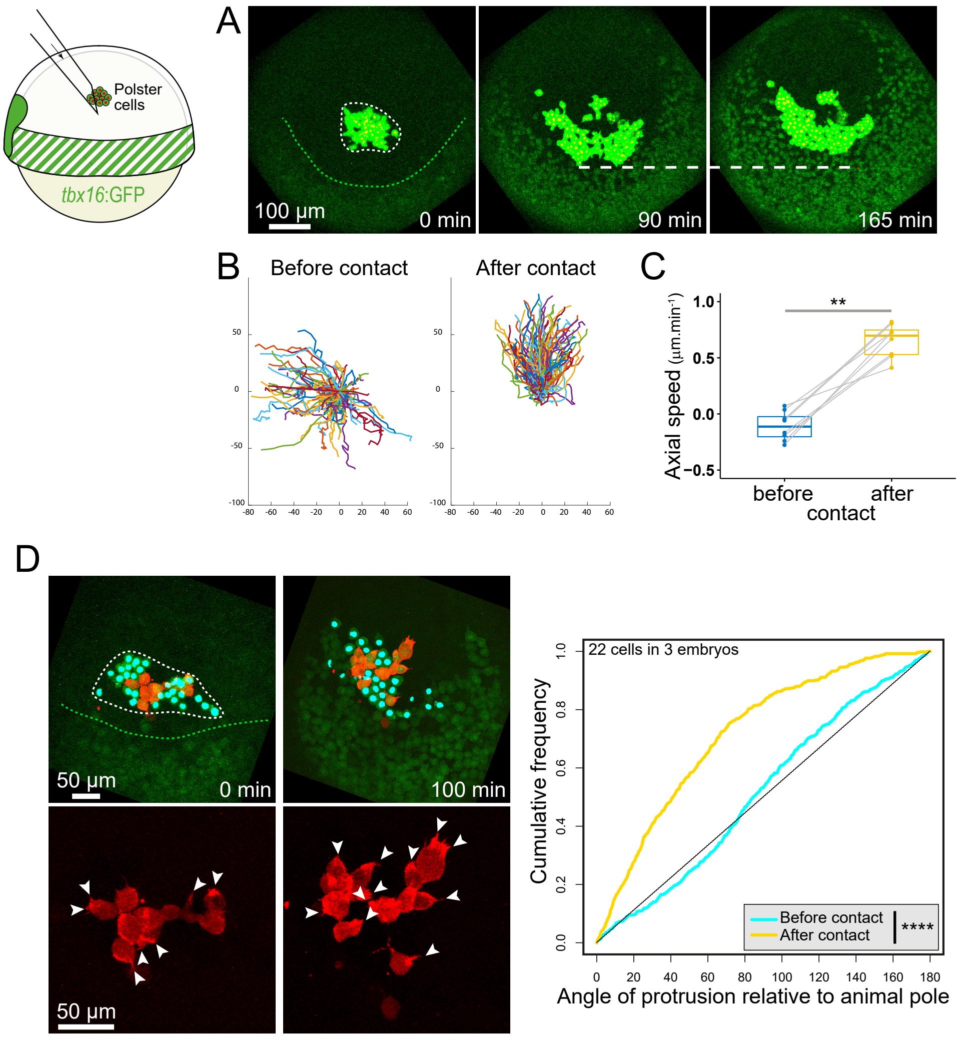


**Supplemental Figure 4. Lateral mesoderm can drive polster cell migration,** related to Figure 4.
(A) H2B-mCherry expressing polster cells (strong green, red nuclei) were transplanted ahead of the lateral mesoderm (Tg(tbx16:GFP); faint green). Thin white line delineates polster cells; green line marks the lateral mesoderm; horizontal line marks the position of the rear of the polster cell group at the moment of contact.
(B) Polster cell trajectories before and after contact with lateral mesoderm of a typical experiment.
(C) Axial speed (n=8 embryos) of transplanted polster cells before and after contact with the lateral mesoderm.
(D) Protrusion orientation of Lifeact-mCherry expressing polster cells (red labelled actin) transplanted along with other polster cells expressing H2B-mCerulean (green with blue nuclei) in front of the lateral mesoderm, quantified before and after contact (22 red cells in 3 embryos).

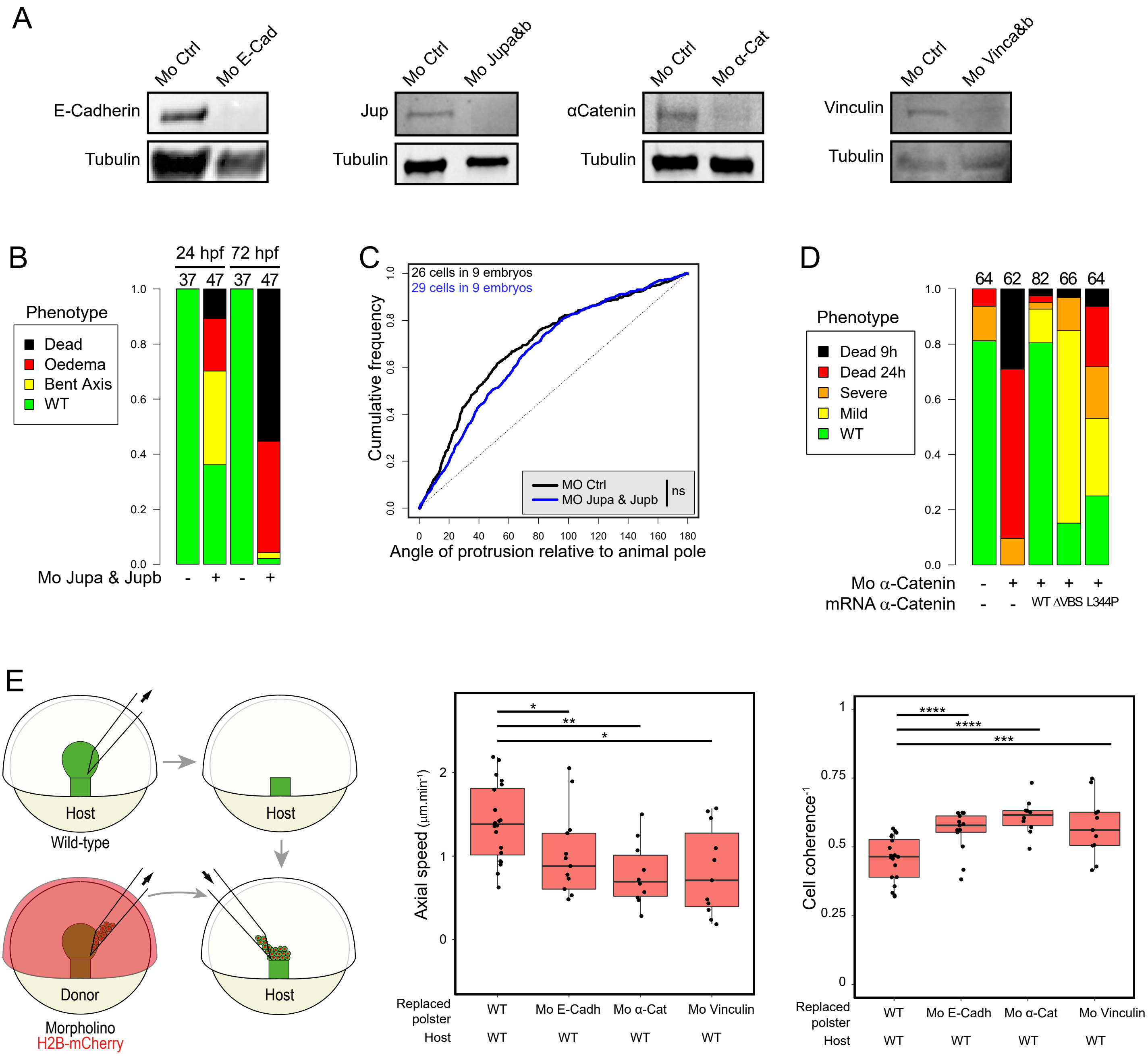

**Supplemental Figure 5. E-Cadherin, α-Catenin and Vinculin are required for polster cell migration,** related to Figure 5.

(A) Western blots with the indicated antibodies, on embryos injected with a control morpholino (Mo Ctrl), or morpholinos targeting e-cadherin (Mo E-Cad), jupa and jupb (Mo Jupa&b), α-catenin (Mo α-Cat), vinculin a & vinculin b (Mo Vinca&b). Each experiment was repeated 4 times.

(B) Phenotypes at 24 and 72 hpf of control uninjected embryos and embryos injected with jupa and jupb morpholinos.

(C) Orientation of protrusions of polster cells injected with Lifeact-mCherry RNAs and a control Morpholino (n=26 in 9 embryos) or Morpholinos targeting jupa and jupb (n=29 cells in 9 embryos), transplanted in a wild-type polster.

(D) Phenotypes at 24 hpf of embryos injected with a control morpholino or Mo α-Catenin, with or without indicated mRNAs. (B,D) Number of analyzed embryos are indicated above each bar.

(E) Replacement of the WT polster by a polster injected with a morpholino targeting E-Cadherin, α-Catenin or Vinculin. Axial speed and coherence-1 (changes in relative distances between pairs of neighbouring cells over time) of polster cells were quantified by tracking cell nuclei.

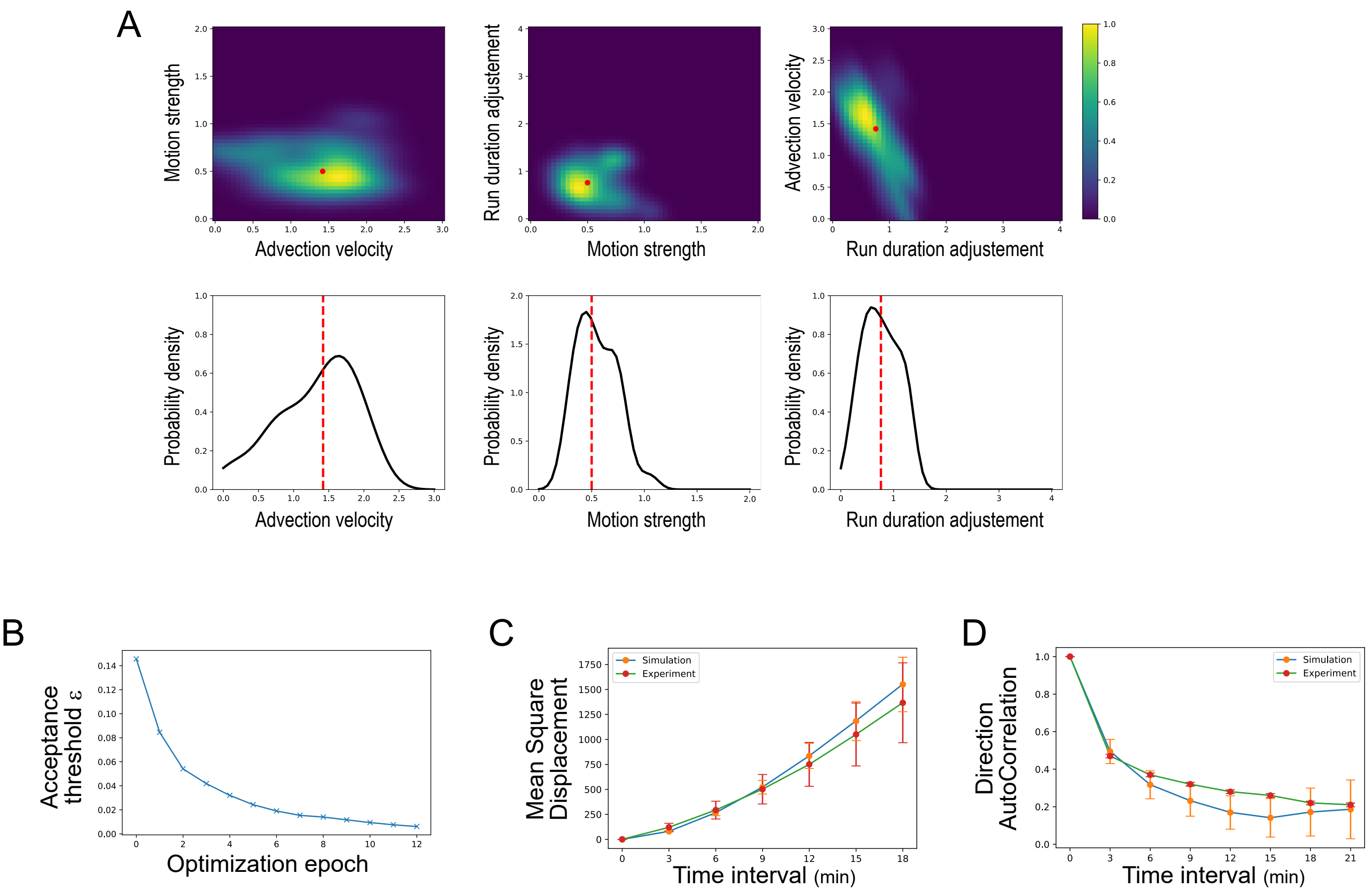


**Supplemental Figure 6. Cellular Potts Model simulations,** related to Figure 7.
(A) Inference of model parameters from experimental data with ABC-SMC optimization using the FitMulticell toolbox. Estimation results are shown as normalized two-parameter (top row) and one-parameter (bottom-row) kernel density estimates of the posterior parameter distributions. The red dot (top row) and red-dashed line (bottom row) represent the weighted median (for details see Methods, Parameter estimation).
(B) Evolution of the acceptance threshold through optimization epochs.
(C) Mean Square Displacement and (D) Differential AutoCorrelation of experimental and simulated data with estimated parameters.

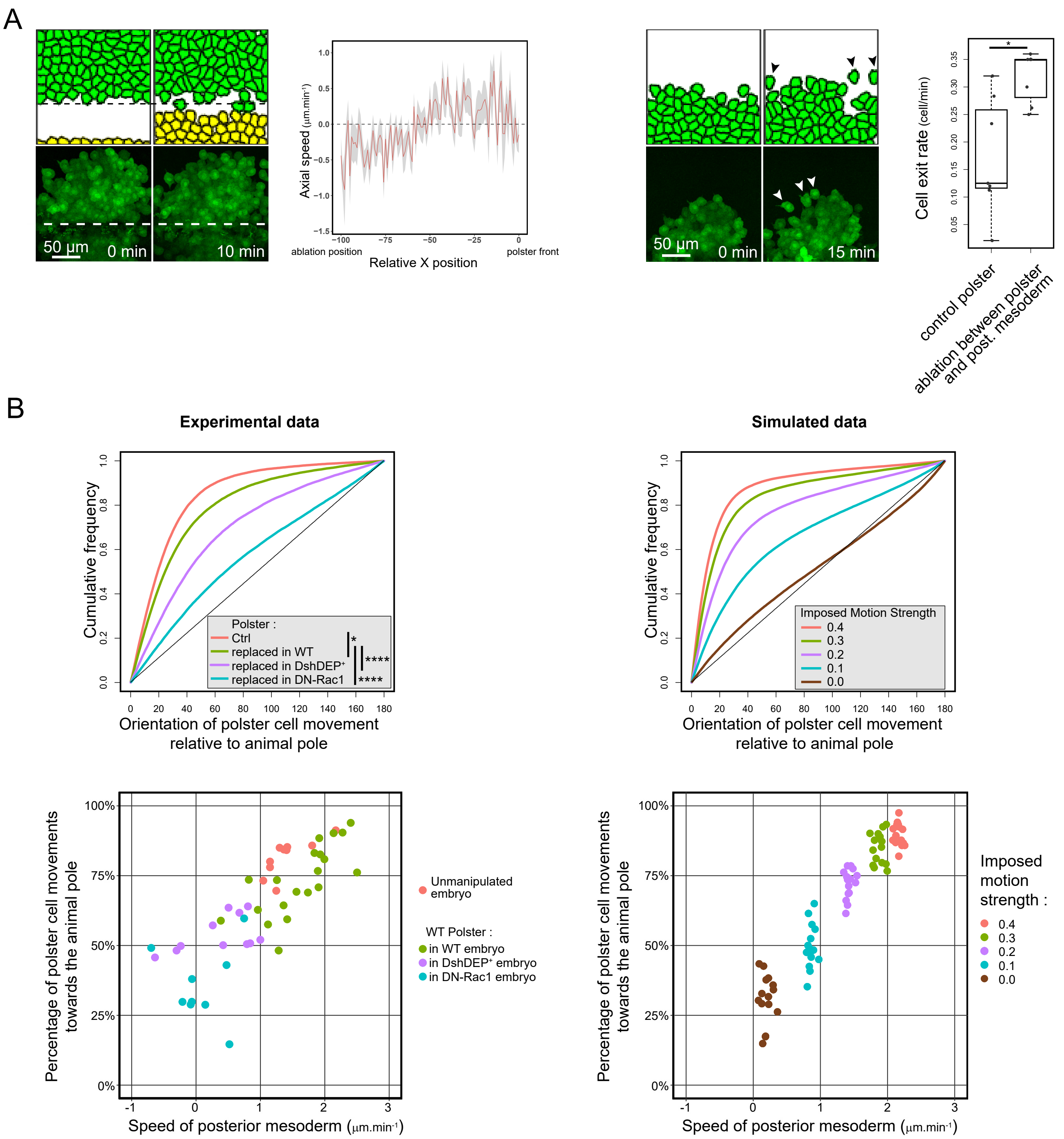


**Supplemental Figure 7. Comparison of simulated and experimental data,** related to Figure 7.
(A) Close-ups from simulated and experimental laser ablations, revealing two features we first noticed in simulations and then identified in experimental data: a backward movement of the cells at the posterior edge of the isolated group and a tendency of the most anterior cells to migrate out of the group. Both features were quantified in experimental data.
(B) Orientation of polster cell movements when speed of following cells (posterior mesoderm) varies. Experimental data are from experiments presented in Figure 4A-D, and correspond to an unmanipulated polster (ctrl), or a wild-type polster transplanted in either a wild-type embryo, a Dsh-DEP+ injected embryo or a DN-Rac1 injected embryo. Simulations were performed with varying speed of posterior mesoderm cells, by modulating the Lagrange multiplier modulating their movement (Imposed Motion Strength). Distribution of orientations are plotted in the different situations as cumulative plots. The percentage of movements oriented towards the animal pole (<45%) is plotted as a function of the measured speed of the axis.
